# Metabolism-Induced Oxidative Stress and DNA Damage Selectively Trigger Genome Instability in Polyploid Cells

**DOI:** 10.1101/480822

**Authors:** Gregory J. Thomson, Claire Hernon, O.P. Nicanor Austriaco, Rebecca S. Shapiro, Peter Belenky, Richard J. Bennett

## Abstract

Understanding the forces impacting genome stability is important for diverse processes such as tumorigenesis and reproductive biology. The pathogenic yeast *Candida albicans* displays unusual genome dynamics in which tetraploid cells, but not diploid cells, become unstable when grown on a glucose-rich ‘pre-sporulation’ medium. Here, we reveal that *C. albicans* polyploid cells are metabolically hyperactive on this medium as evidenced by increased expression of metabolic genes as well as higher rates of fermentation and oxidative respiration. These cells also show elevated levels of reactive oxygen species (ROS), activate the ROS-responsive transcription factor Cap1, and accrue DNA double-strand breaks. Suppression of ROS levels reduced oxidative stress, DNA damage and chromosome instability. These studies reveal how metabolic flux can generate endogenous ROS, triggering DNA damage and genome instability in polyploid, but not diploid, cells. We discuss parallels with metabolism-induced instability in cancer cells and propose that ROS-induced DNA damage could have facilitated ploidy cycling in eukaryotes prior to the evolution of meiosis.

## Introduction

Polyploidy, a state where cells contain more than two copies of their genome, is ubiquitous amongst eukaryotic organisms. In *Arabidopsis*, polyploid cells can arise via gamete fusion or whole genome duplication (Bomblies and Madlung, 2014), and provide advantages such as heterosis, gene redundancy, and enhanced nutrient uptake due to polyploid cells being larger than diploid cells (Chao et al., 2013; Comai, 2005). In humans, polyploidization is known to occur in heart muscle, liver, brain, and blood cells, and is associated both with normal development as well as pathological conditions (Gentric and Desdouets, 2014). Polyploid cells similarly form in uterine muscle tissue as a normal part of pregnancy and are thought to aid in embryonic implantation (Mori et al., 2011; Sroga et al., 2012). Cancer progression is also associated with polyploidization; aneuploid tumors often arise through formation of a tetraploid intermediate and can promote malignancy by disrupting regulators of cell growth (Fujiwara et al., 2005; Storchova and Pellman, 2004; Zack et al., 2013).

Polyploid forms have also been characterized in diverse fungal lineages, including the budding yeast *Saccharomyces cerevisiae* which can exist in ploidy states from 1N to 4N (Zhu et al., 2016). Higher ploidy states can arise via gamete fusion during a conventional sexual cycle or via asexual processes such as mitotic collapse. A characteristic feature of polyploid cells is increased genome instability, with tetraploid cells exhibiting rates of chromosome loss that are orders of magnitude greater than those of diploid cells (Mayer and Aguilera, 1990). Polyploidy and genome instability can facilitate evolution, with higher ploidy states undergoing adaptation faster than diploid or haploid cells due to higher mutation rates with greater fitness effects (Selmecki et al., 2015). Ploidy shifts in *S. cerevisiae* cells are induced by a variety of environmental stresses, including culture in the presence of high salt or ethanol (Gerstein et al., 2006; Voordeckers et al., 2015).

Ploidy variation has also been described in the opportunistic fungal pathogen *C. albicans*, a commensal species that can seed both debilitating mucosal and life-threatening systemic infections. Candidiasis is a major problem in the hospital setting and is exacerbated by the prevalence of drug-resistant *C. albicans* strains (Ksiezopolska and Gabaldon, 2018). Natural *C. albicans* isolates are diploid although rare haploid forms have been isolated in the laboratory, albeit with reduced fitness (Hickman et al., 2013). As in *S. cerevisiae*, ploidy shifts can enable adaptation; diploid *C. albicans* cells exposed to the antifungal drug fluconazole form a transient tetraploid state that rapidly generates drug-resistant aneuploid progeny via aberrant mitotic divisions (Harrison et al. 2014).

Tetraploid *C. albicans* cells are also formed by fusion of diploid cells during mating (Bennett and Johnson, 2003; Hull et al., 2000; Magee and Magee, 2000). Sexual reproduction in *C. albicans* is unusual in that cells must undergo an epigenetic switch from the sterile ‘white’ state to the mating-competent ‘opaque’ state (Bennett, 2015; Lockhart et al., 2002; Miller and Johnson, 2002). However, rather than undergoing a conventional meiosis, tetraploid cells return to the diploid (or aneuploid) state via aberrant mitotic divisions, a process termed “concerted chromosome loss” (CCL) (Bennett and Johnson, 2003; Forche et al., 2008; Hickman et al., 2015). Growth on *S. cerevisiae ‘*pre-sporulation’ (PRE-SPO) medium at 37°C induces CCL in tetraploid cells, while diploid cells remain stable under these culture conditions. In addition to genome instability, tetraploid cells undergo extensive cell death during growth on PRE-SPO medium whereas diploid cells remain viable. However, how PRE-SPO medium induces polyploid-specific responses in *C. albicans* cells has not been determined.

In this work, we reveal the mechanism by which a shift in metabolism triggers genome instability in polyploid, but not diploid, cells. Transcriptional and metabolic profiling establish that *C. albicans* cells grown on glucose-rich PRE-SPO medium are metabolically hyperactive, including increased rates of both fermentation and oxidative respiration. Critically, cellular metabolism is further upregulated in cells of higher ploidy in line with their increased cell size. The consequence of increased metabolic flux is the production of reactive oxygen species (ROS) and consequent DNA damage that is the primary cause of genome instability in polyploid cells. These studies therefore establish direct links between metabolic flux, ROS, oxidative stress and DNA damage, and reveal that this mechanism can selectively impact genome integrity in polyploid cells. We also hypothesize that metabolism-induced genome instability was responsible for a primitive ploidy cycle in early eukaryotes prior to the emergence of a conventional meiosis.

## Results

### Polyploid *C. albicans* cells undergo CCL on PRE-SPO medium

A conventional meiosis has not been observed in *C. albicans* despite its genome containing numerous conserved meiosis genes (Butler et al., 2009; Tzung et al., 2001). Instead, tetraploid mating products reduce their ploidy via a parasexual program of CCL, during which tetraploid cells return to a diploid or aneuploid state (Bennett and Johnson, 2003; Forche et al., 2008). CCL can be induced by incubation of tetraploid *C. albicans* cells on *S. cerevisiae* PRE-SPO medium at 37°C (Bennett and Johnson, 2003). Tetraploid cells experience extensive cell death during growth on PRE-SPO medium at 37°C whereas most diploid cells remain viable under these culture conditions (Figure 1A), as previously reported (Bennett and Johnson, 2003). Tetraploid cells grown on PRE-SPO medium also exhibit erratic microtubule and nuclear dynamics, indicative of aberrant nuclear divisions on this medium (Figure 1B).

**Figure 1.**
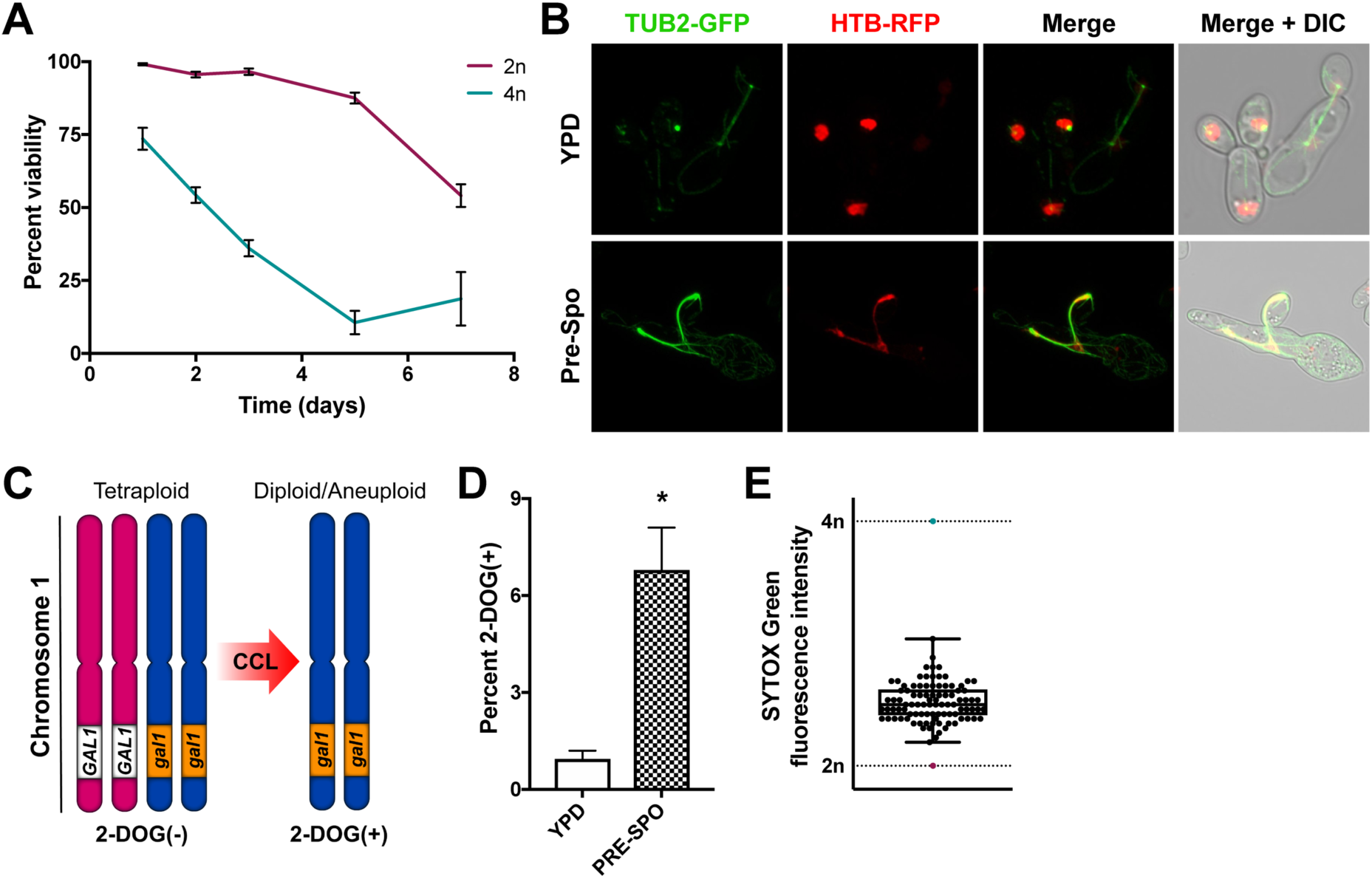
*C. albicans* tetraploid cells exhibit genome instability on PRE-SPO medium. **(A)** Viability of diploid (SC5314) and tetraploid (RBY18) cells cultured on PRE-SPO medium at 37°C. (**B)** A tetraploid β-tubulin (TUB2)-GFP, histone H2B (HTB)-RFP stain was cultured on YPD or PRE-SPO medium at 30°C or 37°C, respectively, for 24 h and imaged. Cell images indicate maximum intensity Z projections for GFP and RFP channels, and merged images of the GFP/RFP and GFP/RFP/DIC channels. (**C)** Schematic illustrating assay to monitor for loss of *GAL1* in tetraploid strain RBY18. Strain RBY18 is heterozygous for the *GAL1* gene (*GAL1/GAL1/gal1/gal1*) and only cells that have undergone loss of both wild type copies of *GAL1* can grow on 2-DOG medium. (**D)** Tetraploid strain RBY18 (*GAL1/GAL1/gal1/gal1*) was cultured on YPD and PRE-SPO media at 37°C for seven days and plated onto 2-DOG medium to monitor for loss of the *GAL1* gene. (**E)** 94 2-DOG^R^ progeny arising from RBY18 on PRE-SPO medium were DNA stained and their ploidy determined via flow cytometry. Upper and lower whiskers represent the maximum and minimum values from the analyzed progeny.

To monitor genome stability in tetraploid cells, we used a strain, RBY18, that is heterozygous for *GAL1* on chromosome 1 (*GAL1/GAL1/gal1/gal1*) (Figure 1C). Only strains that have lost *GAL1* function are viable on 2-deoxygalactose (2-DOG) medium (Gorman et al., 1992), providing a selection for cells that have undergone a reduction in ploidy and lost the two wildtype *GAL1* alleles (Bennett and Johnson, 2003). When RBY18 was cultured on PRE-SPO medium at 37°C for 7 days, approximately 7% of cells lost *GAL1* and became 2-DOG^R^ (Figure 1D). Flow cytometric analysis of DNA content in 2-DOG^R^ colonies indicated that tetraploid (4N) cells had undergone a reduction in ploidy and presented with a DNA content between 2N and 3N (Figure 1E).

### Transcriptional profiling of diploid and tetraploid cells on YPD and PRE-SPO media

RNA-sequencing (RNA-seq) was performed to determine gene expression of *C. albicans* cells grown on standard YPD medium (at 30°C) and on PRE-SPO medium (at 37°C). YPD cultures were performed at the lower temperature as extensive filamentation was observed during growth on YPD, but not PRE-SPO, medium at 37°C. Both diploid and tetraploid cells were analyzed on these media at 3, 6, 12, and 24 h time points.

RNA-seq analysis revealed that diploid and tetraploid cells showed similar gene expression profiles on YPD and PRE-SPO media, despite differences in cell survival and genome stability on the two media (Figure 2A). Notably, however, gene ontology (GO) analysis found that processes related to DNA double-strand break (DSB) formation, DNA repair, DNA recombination, and meiosis were significantly enriched among differentially expressed genes between cells grown on YPD and PRE-SPO medium at 24 h, as were genes involved in stress and metabolism (Figure 2B). Analysis of gene expression in cells grown on PRE-SPO medium revealed that most meiosis-specific genes were not induced except for genes involved in meiotic recombination (Supplemental Figure 1). We also conducted GO term analysis on differentially expressed genes in diploid v. tetraploid cells grown on PRE-SPO medium at each time point. At 3 and 6 h, nicotinamide binding and acetate metabolism genes were enriched among differentially expressed genes (Supplemental Tables 1,2), while at 12 and 24 h, DNA replication, regulation of helicase activity, and DNA binding genes were enriched among differentially expressed genes (Supplemental Tables 3,4). These results implicated metabolism and DNA damage as being relevant to the distinct behavior of tetraploid cells grown on PRE-SPO versus YPD medium.

**Figure 2.**
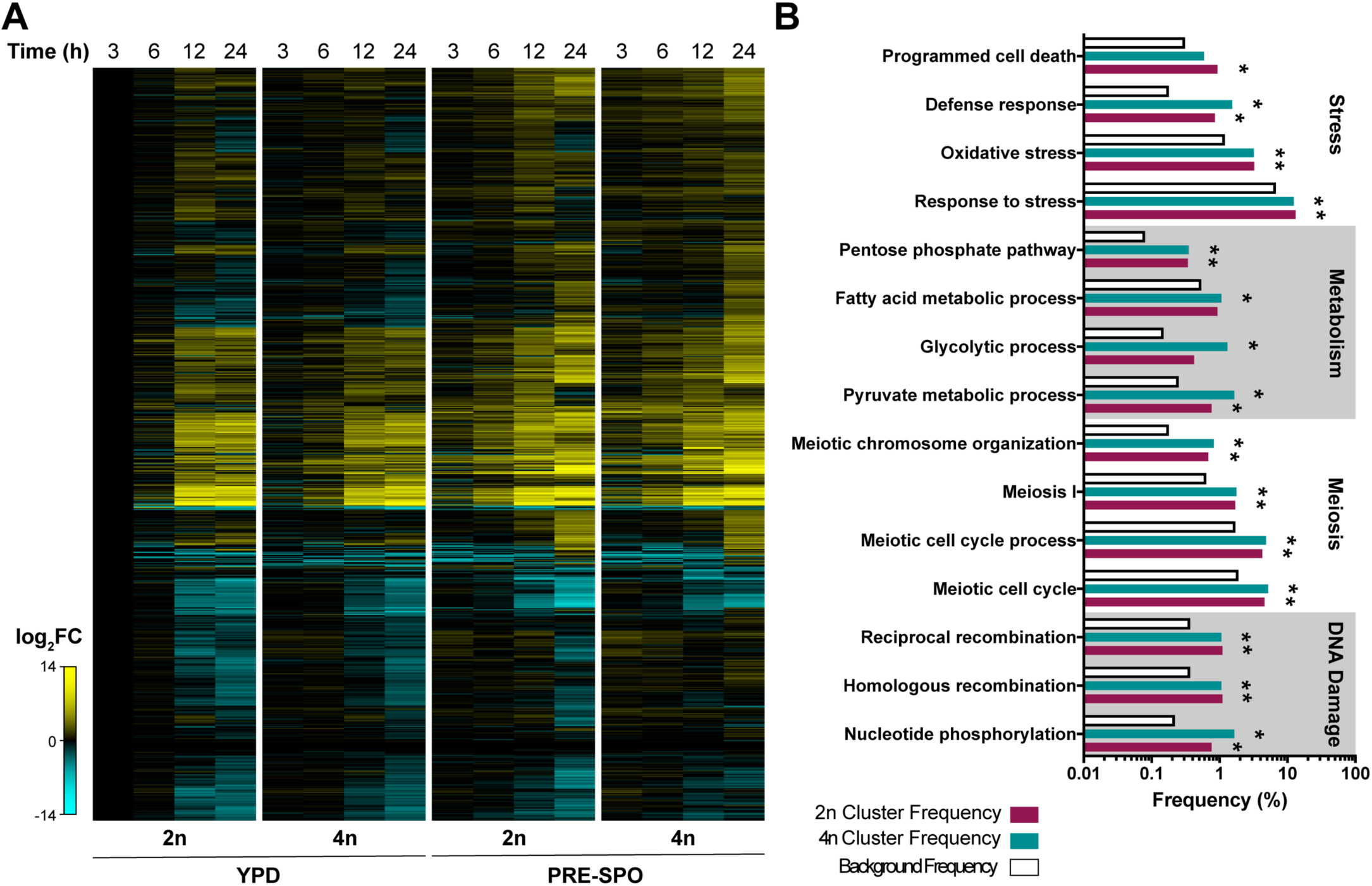
Transcriptional profiling of *C. albicans* diploid and tetraploid cells on YPD and PRE-SPO medium. **(A)** Heatmap of RNA-Seq data showing global gene expression changes in diploid (SC5314) and tetraploid (RBY18) cells on YPD or PRE-SPO medium over 24 h. (**B)** Results of gene ontology (GO) term enrichment analysis of differentially expressed genes between YPD and PRE-SPO media at 24 h in diploid and tetraploid cells. Background frequency represents the total number of genes annotated to each GO term in the *C. albicans* genome divided by the total number of genes in the genome. (**\***) denotes a statistically significant enrichment (p<0.05) of genes annotated to the indicated GO term.

Inspection of metabolic gene expression revealed that glycolytic genes were induced earlier during growth on PRE-SPO than on YPD medium in both diploid and tetraploid cells (compare 6 h time points, Figure 3A). In addition, oxidative stress-associated genes, including those encoding the Sod3 superoxide dismutase and the Ddr48 DNA damage response factor, were activated at earlier time points on PRE-SPO than on YPD medium. These results indicate that growth on PRE-SPO medium results in enhanced metabolic activity and activation of oxidative stress response genes.

**Figure 3.**
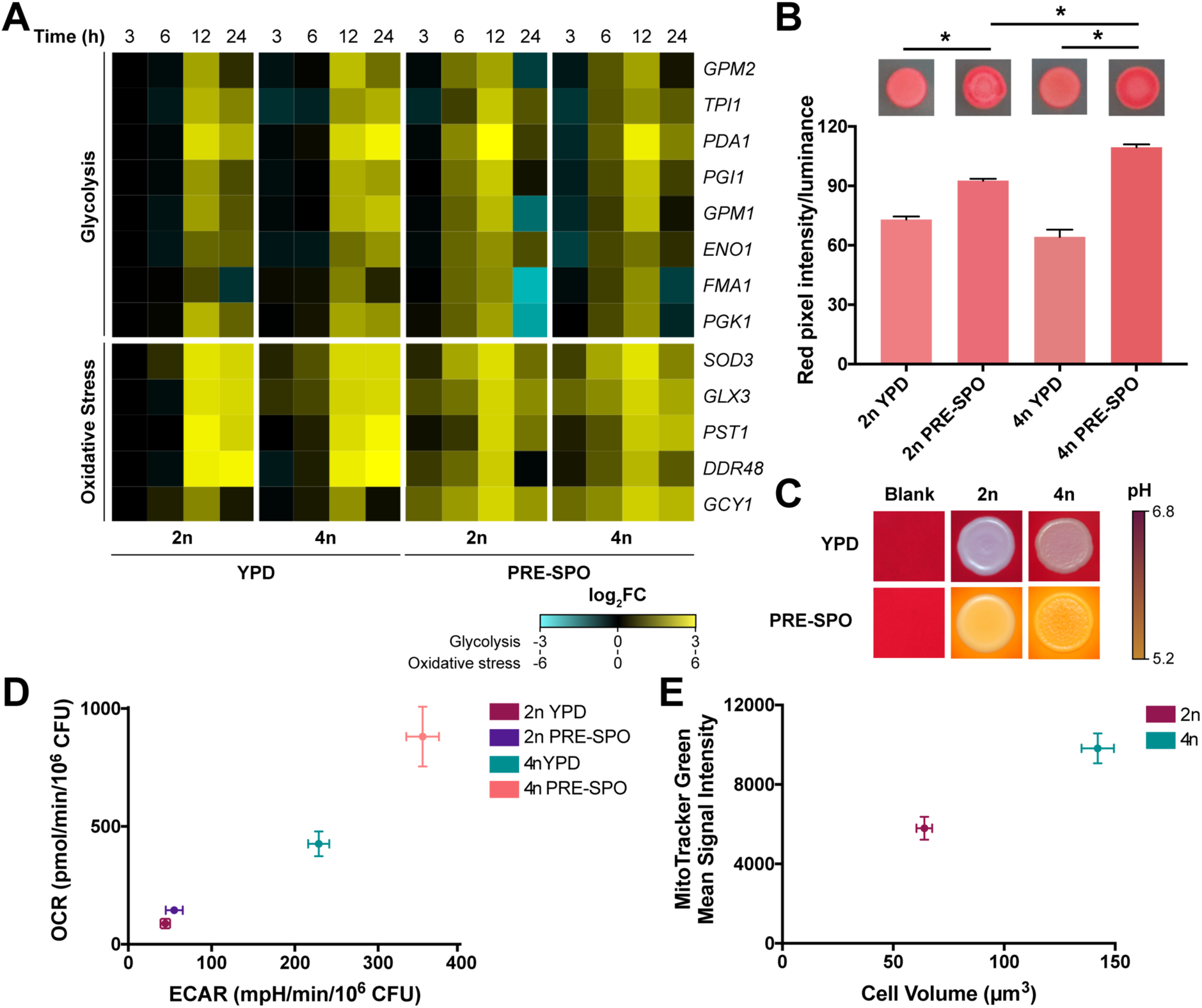
Analysis of metabolic and oxidative stress responses in diploid and tetraploid cells grown on YPD and PRE-SPO medium. **(A)** Heatmap of RNA-seq data for glycolysis and oxidative stress genes in diploid and tetraploid cells grown on YPD and PRE-SPO media over 24 h. (**B)** TTC assay results in diploid and tetraploid cells grown on YPD and PRE-SPO media. Cells were grown for 24 h at which point a TTC overlay solution was applied and red color allowed to develop for 15 min. Calculated values represent the total amount of reddening in each colony. (**\***) denotes a significant difference (p<0.05). (**C)** Diploid and tetraploid strains were spot inoculated onto YPD and PRE-SPO media supplemented with the pH indicator bromocresol purple and allowed to grow for 24 h. (**D)** Analysis of metabolic flux in diploid and tetraploid cells grown on YPD and PRE-SPO media for 24 h at 37°C using a Seahorse instrument. OCR values represent rates of aerobic respiration while ECAR values represent rates of fermentation. (**E)** Cell size and mean MitoTracker Green signal intensity in diploid and tetraploid cells grown on YPD medium for 24 h at 30°C.

### *C. albicans* tetraploid cells are metabolically hyperactive on PRE-SPO medium

Given that *C. albicans* cells show more rapid activation of glycolytic and oxidative stress genes on PRE-SPO than on YPD medium, we compared the metabolic activity of diploid and tetraploid cells on the two media using a 2,3,5-triphenyltetrazolium chloride (TTC) overlay technique. TTC is a white compound that passively diffuses through plasma membranes and interacts with mitochondria where it is converted to red 1,3,5-triphenylformazan (TPF). TTC overlay assays are commonly used to assess activity of the electron transport chain in microbial populations (Morales et al., 2013; Ogur et al., 1957; Rich et al., 2001). Equivalent numbers of diploid and tetraploid *C. albicans* cells were spotted onto YPD and PRE-SPO media and grown for 24 h at 30°C and 37°C, respectively, at which point TTC was applied to each plate. We found that both diploid and tetraploid cells exhibited greater reddening on PRE-SPO medium than on YPD medium, with tetraploid cells also generating more color than diploid cells suggestive of heightened respiratory activity (Figure 3B).

Next, we examined media acidification which indicates the formation of acidic fermentation end-products such as acetate (Morales et al., 2013). YPD and PRE-SPO media were supplemented with the pH indicator bromocresol purple, which is yellow at pH values below 5.2 and purple at pH values above 6.8. Equal numbers of diploid and tetraploid cells were inoculated onto YPD and PRE-SPO media and grown for 24 h at 30°C and 37°C, respectively. Diploid and tetraploid cells both acidified PRE-SPO medium whereas the pH was largely unchanged with cells grown on YPD medium (Figure 3C), suggesting increased production of acidic fermentation end-products during growth on PRE-SPO medium.

To directly characterize the metabolic activities of diploid and tetraploid cells, aerobic respiratory and fermentative rates were assayed by measuring the oxygen consumption rate (OCR) and extracellular acidification rate (ECAR), respectively. *C. albicans* cells were grown on YPD and PRE-SPO media (both for 24 h at 37°C) and transferred to a Seahorse instrument to measure metabolic rates. We found that cells of higher ploidy exhibited significantly higher metabolic activities; both OCR and ECAR were ∼5-fold higher on YPD medium and ∼6 fold higher on PRE-SPO medium in tetraploid cells compared to diploid cells (Figure 3D). We also found that *C. albicans* cells grown on PRE-SPO medium exhibited a significantly higher OCR and ECAR relative to the same cells grown on YPD medium. Tetraploid cells showed a 1.5-fold increase in OCR and a 2.1-fold increase in ECAR when grown on PRE-SPO medium relative to YPD medium, and diploid cells showed similar fold increases in metabolic rates on PRE-SPO medium (Figure 3D). Together, these results establish the impact of both cell ploidy and culture media on *C. albicans* metabolism; growth on PRE-SPO medium enhances metabolic activity relative to growth on YPD medium, and tetraploid cells are also considerably more metabolically active than diploid cells.

Given that tetraploid cells exhibited a higher OCR than diploid cells, we sought to probe the relationship between ploidy and mitochondrial content. Diploid and tetraploid cells were grown on YPD medium for 24 h at 30°C and stained with the mitochondrial marker MitoTracker Green. Cells were then analyzed by flow cytometry to determine mitochondrial content and by microscopy to examine cell size. Tetraploid cells had a cell volume ∼2.2-fold greater than diploid cells and this correlated with an increased MitoTracker Green signal indicating an increased mitochondrial content. These results demonstrate that *C. albicans* tetraploid cells are more metabolically active than diploid cells, which is likely a consequence, at least in part, of their larger cell size and increased mitochondrial content.

### *C. albicans* cells experience elevated oxidative stress on PRE-SPO medium

Transcriptional profiling showed that oxidative stress genes were upregulated in *C. albicans* cells grown on PRE-SPO medium. We therefore examined oxidative stress responses on YPD and PRE-SPO media. To determine if reactive oxygen species (ROS) were present, diploid and tetraploid cells were grown for 24 h on YPD and PRE-SPO media at 30°C and 37°C, respectively, and stained with CellROX Green, a membrane permeant dye that exhibits bright fluorescence upon oxidation by ROS and binding to DNA. A CellROX Green signal was absent from cells grown on YPD medium, whereas both diploid and tetraploid cells grown on PRE-SPO medium stained positively indicating that ROS are generated during growth on this medium (Figure 4A, Supplemental Figure 2A). Flow cytometric analysis of CellROX Green-stained cells corroborated cell microscopic images, with an increased fluorescence signal in diploid and tetraploid cells on PRE-SPO medium relative to YPD medium (Figure 4B). Notably, tetraploid cells on PRE-SPO medium exhibited the strongest signal, with 2.0-fold higher fluorescence levels than diploid cells on this medium.

**Figure 4.**
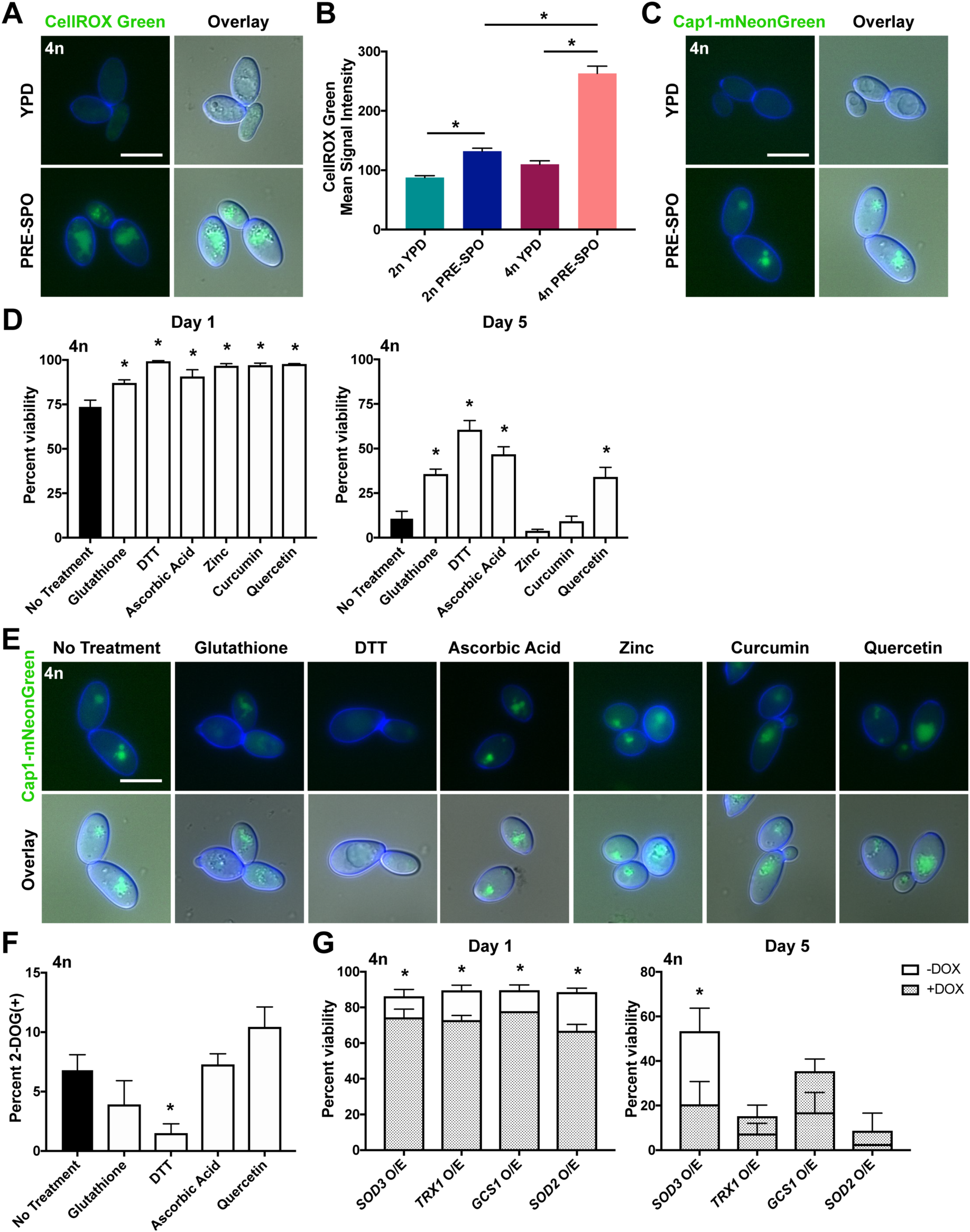
*C. albicans* cells accumulate ROS and activate oxidative stress responses when cultured on PRE-SPO medium. **(A)** Tetraploid cells were grown on YPD or PRE-SPO medium at 30°C or 37°C, respectively, for 24 h and stained with CellROX Green. Cell images indicate calcofluor white staining (cell wall; blue), GFP, and a merged image of GFP/DAPI/DIC channels. Scale bar = 10 µm. (**B)** Diploid and tetraploid cells were grown on YPD or PRE-SPO medium at 30°C or 37°C, respectively, for 24 h, stained with CellROX Green, and signal analyzed via flow cytometry. (*) denotes a significant difference (p<0.05). (**C)** Tetraploid cells containing a fluorescently labeled version of the transcription factor Cap1 (Cap1-mNeonGreen) were grown on YPD or PRE-SPO medium at 30°C or 37°C, respectively, for 24 h. Cell images indicate calcofluor white staining (cell wall; blue), GFP, and a merged image of GFP/DAPI/DIC channels. Scale bar = 10 µm. (**D)** Analysis of cell viability after 1d (top panel) or 5d (bottom panel) of tetraploid cells cultured on PRE-SPO medium supplemented with different antioxidants. (*) denotes a significant difference between cells with and without the indicated antioxidant (p<0.05). (**E)** Tetraploid cells containing Cap1-mNeonGreen were grown on PRE-SPO medium supplemented with antioxidants for 24 h at 37°C. Cell images indicate calcofluor white staining (cell wall; blue), GFP, and a merged image of GFP/DAPI/DIC channels. Scale bar = 10 µm. (**F)** Assay to monitor for loss of *GAL1* from tetraploid strain RBY18 after growth on untreated PRE-SPO medium or PRE-SPO medium supplemented with antioxidants at 37°C for 7 d. (**\***) denotes a significant difference (p<0.05). (**G)** Analysis of cell viability after 1 d (left panel) or 5 d (right panel) in tetraploid cells overexpressing the indicated genes. Cells were grown for 24 h on PRE-SPO medium in the presence (gene OFF) and absence (gene ON) of doxycycline. (**\***) denotes a significant difference between viability in the + and – DOX conditions (p<0.05).

Previous work showed that *C. albicans* produces ROS during hyphal morphogenesis (Rossi et al., 2017). Consistent with this finding, cells grown on YPD medium at 37°C underwent filamentation and stained positively with CellROX Green (Supplemental Figure 3A). However, a fluorescence signal was absent in cells grown on SCD medium at 37°C (Figure 7A), a growth condition where efficient filamentation does not occur, indicating that growth at 37°C is not sufficient for production of ROS.

We also examined oxidative stress responses using a fluorescently tagged Cap1 protein, a transcription factor that regulates multiple oxidative stress response pathways in *C. albicans* including antioxidant defense, protein degradation and drug resistance (Wang et al., 2006). Cap1 is distributed homogenously between the nucleus and the cytoplasm under non-stress conditions, but accumulates in the nucleus in response to oxidative stress to activate antioxidant gene expression (Patterson et al., 2013). To assay for Cap1 activation, diploid and tetraploid strains were generated expressing fluorescently labeled Cap1-mNeonGreen. These reporter strains were grown for 24 h on YPD and PRE-SPO media at 30°C and 37°C, respectively, and imaged to examine Cap1 localization. Diploid and tetraploid cells grown on PRE-SPO medium exhibited a clear nuclear Cap1 signal whereas cells grown on YPD medium exhibited only a diffuse signal (Figure 4C, Supplemental Figure 2B). The increased temperature alone was not responsible for the observed phenotypes on PRE-SPO medium, as tetraploid cells grown on YPD medium at 37°C for 24 h did not exhibit a nuclear Cap1 signal (Supplemental Figure 3B). These results establish that both diploid and tetraploid cells experience elevated levels of oxidative stress when grown on PRE-SPO medium.

To determine if oxidative stress contributes to decreased cell viability in tetraploid cells on PRE-SPO medium, this medium was supplemented with the antioxidants quercetin, curcumin, ascorbic acid, zinc sulfate, glutathione or dithiothreitol (DTT). All of these antioxidants significantly increased viability after 1 day of growth on PRE-SPO medium, with glutathione, DTT, quercetin and ascorbic acid providing protection out to 5 days of growth (Figure 4D). Glutathione and DTT also abrogated the nuclear Cap1 signal indicating these chemicals protected cells from oxidative stress (Figure 4E). Antioxidants that provided a long-term rescue in viability were also tested for their ability to mitigate genome instability in tetraploid cells grown on PRE-SPO medium. DTT significantly reduced genome instability whereas glutathione produced a partial (but not significant) reduction in genome instability (Figure 4F).

We also assessed the ability of endogenous enzymatic activities to protect against tetraploid cell death on PRE-SPO medium. Tetraploid strains were constructed containing a doxycycline repressible ‘Tet-OFF’ cassette to regulate overexpression (O/E) of several antioxidant genes including *SOD2*, encoding the mitochondrial manganese-containing superoxide dismutase, *SOD3*, encoding the cytosolic manganese-containing superoxide dismutase, *TRX1*, encoding the major *C. albicans* thioredoxin, and *GCS1*, encoding the enzyme that catalyzes the rate-limiting step in glutathione production. Forced expression of each of the antioxidant enzymes provided a short-term rescue in viability after 1 day of growth on PRE-SPO medium, and the *SOD3 O/E* strain provided a long-term rescue over 5 days of growth on this medium (Figure 4G). Taken together, these results establish that oxidative stress is a major contributor to both tetraploid cell death and chromosome instability on PRE-SPO medium.

### DNA double-strand breaks are generated during growth on PRE-SPO medium

A key mechanism by which ROS induces cellular toxicity in multiple cell types is through structural changes to DNA including base modifications, DNA cleavage events, and DNA-protein crosslinks (Jena, 2012). Given that detectable levels of ROS accrue in cells during growth on PRE-SPO medium and that these cells exhibit activation of oxidative stress responses, we examined whether cells exhibit formation of DNA double-strand breaks (DSBs).

To track formation of DSBs, we generated a reporter assay using the viral Gam protein that binds irreversibly to sites of DSBs. Fluorescent labeling of Gam has been shown to enable the visualization of DSBs in live mammalian and bacterial cells via fluorescence microscopy (Belenky et al., 2015; Shee et al., 2013). Diploid and tetraploid *C. albicans* strains containing a doxycycline-inducible (Tet-ON) Gam-GFP construct were grown for 24 h on YPD and PRE-SPO media at 30°C and 37°C, respectively, and examined by fluorescence microscopy and flow cytometry. An intense nuclear signal was detected in diploid and tetraploid cells cultured on PRE-SPO medium, whereas no signal was detected on YPD medium (Figure 5A, Supplemental Figure 2C). This finding was corroborated by flow cytometry, where approximately 26% of tetraploid cells became Gam-GFP positive after 24 h on PRE-SPO medium in the presence of doxycycline (Figure 5B). The elevated growth temperature was not solely responsible for increased DNA damage on PRE-SPO medium as tetraploid cells cultured on YPD medium at 37°C did not exhibit a nuclear GAM-GFP signal and the percent of Gam-GFP positive cells increased minimally, albeit significantly, relative to growth on YPD medium at 30°C (Supplemental Figure 3C,D).

**Figure 5.**
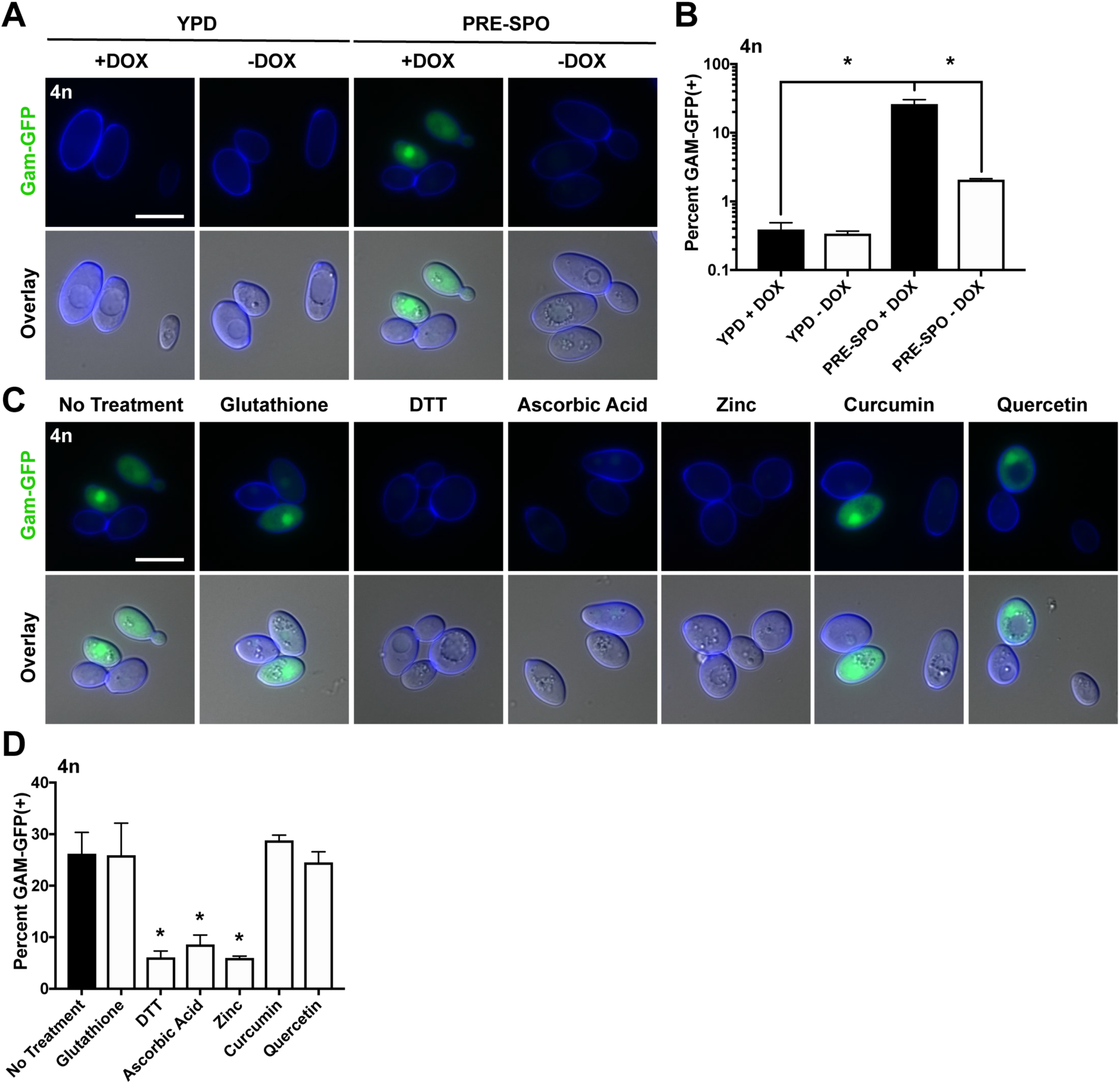
Oxidative stress contributes to DNA double-strand break (DSB) formation during growth on PRE-SPO medium. **(A)** A tetraploid strain expressing a Tet-ON Gam-GFP reporter was cultured on YPD or PRE-SPO medium at 30°C or 37°C, respectively, in the presence (Gam-GFP ON) or absence (Gam-GFP OFF) of doxycycline for 24 h. Cell images indicate calcofluor white staining (cell wall; blue), GFP, and a merged image of GFP/DAPI/DIC channels. Scale bar = 10 µm. (**B)** Flow cytometric analysis of the tetraploid Gam-GFP reporter strain grown on YPD or PRE-SPO medium at 30°C or 37°C, respectively, in the presence or absence of doxycycline for 24 h. (**\***) indicates a significant difference (p<0.05). (**C)** Tetraploid cells expressing a Gam-GFP reporter were cultured on PRE-SPO medium supplemented with the indicated antioxidants at 37°C for 24 h and analyzed by fluorescence microscopy. Cell images indicate calcofluor white staining (cell wall; blue), GFP, and a merged image of GFP/DAPI/DIC channels. Scale bar = 10 µm. (**D)** Flow cytometric analysis of the tetraploid Gam-GFP reporter strain grown on PRE-SPO medium supplemented with antioxidants at 37°C for 24 h. (**\***) denotes a significant difference between untreated PRE-SPO medium and PRE-SPO medium supplemented with the indicated antioxidant (p<0.05).

We next examined if protection from oxidative stress would reduce DSB formation in tetraploid cells grown on PRE-SPO medium. The tetraploid Gam-GFP reporter strain was cultured with and without antioxidants on PRE-SPO medium for 24 h and cells again examined using fluorescence microscopy and flow cytometry. Interestingly, all water-soluble antioxidants except glutathione reduced the formation of DSBs, whereas the fat-soluble antioxidants quercetin and curcumin did not prevent DSB formation (Figure 5C,D).

These results demonstrate that *C. albicans* tetraploid cells experience extensive DSB formation when cultured on PRE-SPO medium, and that chemical agents that diminish ROS levels reduce these high levels of DNA damage.

### Tetraploid cells exposed to DNA damaging agents also undergo CCL

DSBs can present a highly destabilizing form of DNA damage (Jackson and Bartek, 2009; Mehta and Haber, 2014) and we therefore directly tested the effect of DSBs on *C. albicans* cells. Tetraploid cells were exposed to the DNA-damaging agents hydroxyurea (HU) and methyl methanesulfonate (MMS) and the Gam-GFP reporter strain used to evaluate DSB formation. We observed DSB formation in tetraploid cells exposed to both DNA-damaging compounds (Figure 6A,B). Moreover, when assayed for sensitivity to MMS and HU at 30°C and 37°C, tetraploid cells appeared more sensitive to both compounds compared to diploid cells, and this was particularly evident at 37°C (Figure 6C, Supplemental Figure 4).

**Figure 6.**
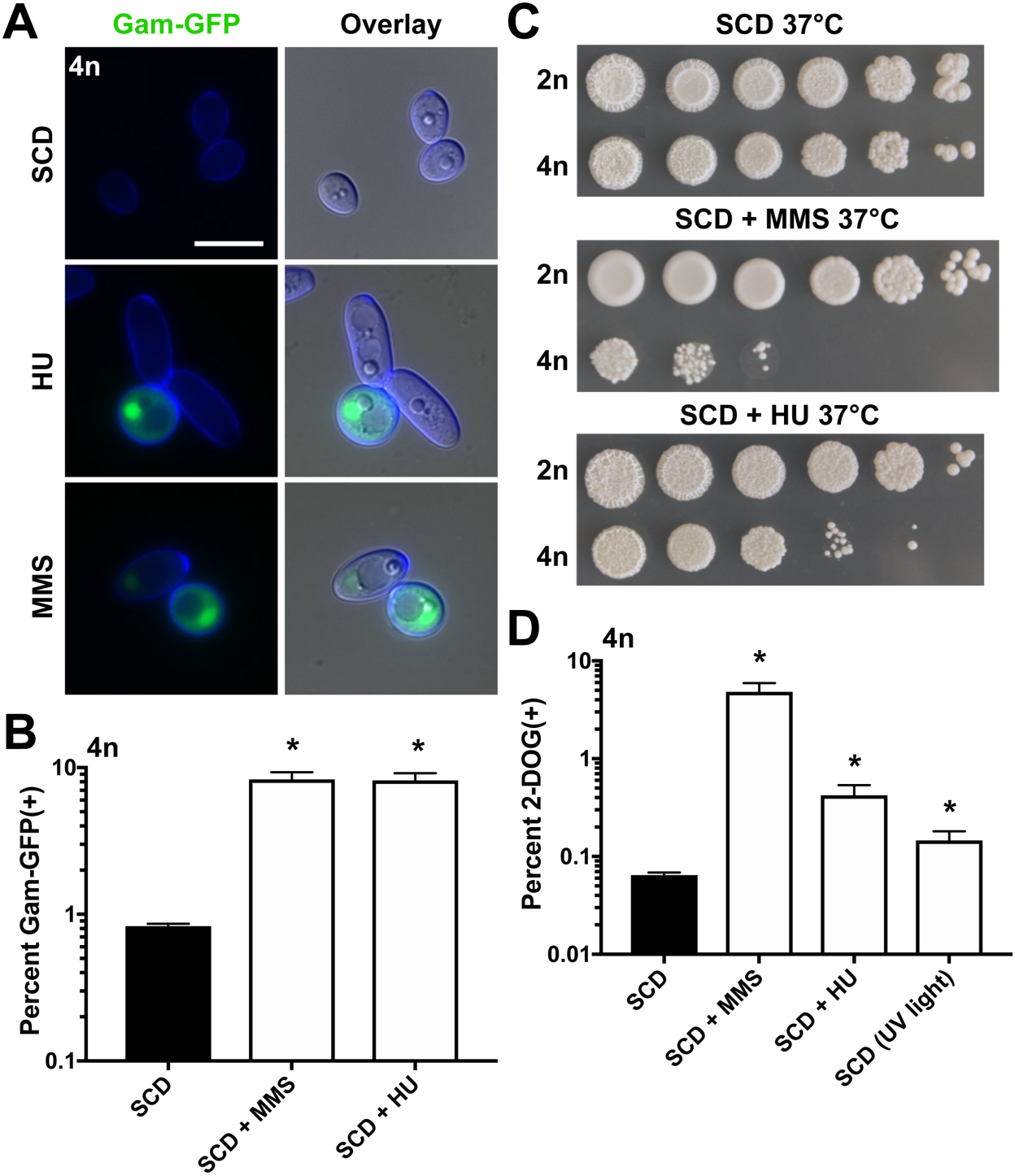
DNA damage induces CCL in tetraploid cells. **(A)** Tetraploid cells expressing a Gam-GFP reporter were cultured on SCD medium supplemented with the DNA damaging agents HU (20mM) or MMS (0.01%) at 37°C for 24 h and analyzed via fluorescence microscopy. Cell images indicate calcofluor white staining (cell wall; blue), GFP, and a merged image of GFP/DAPI/DIC channels. Scale bar = 10 µm. (**B)** Flow cytometric analysis of the tetraploid Gam-GFP reporter strain grown on SCD medium supplemented with 20mM HU or 0.01% MMS at 37°C for 24 h. (**\***) denotes a significant difference between untreated SCD medium and that supplemented with the indicated agent (p<0.05). (**C)** 10-fold dilutions of diploid and tetraploid cells were spot inoculated onto SCD medium and SCD medium containing the DNA-damaging agents HU (20mM) or MMS (0.01%). Cells were allowed to grow for 48 h at 37°C and plates imaged. (**D)** 2-DOG assay to monitor for loss of *GAL1* from tetraploid strain RBY18 (*gal1/gal1/GAL1/GAL1*) after culture on SCD medium supplemented with the DNA-damaging agents HU (20mM) or MMS (0.01%), or exposed to UV light for five seconds a day at 37°C for 7 days. (**\***) denotes a significant difference (p<0.05).

To examine if DNA damage can induce chromosomal instability in *C. albicans* tetraploid cells, the tetraploid RBY18 strain was exposed to HU, MMS, or ultraviolet (UV) radiation and cells assayed for those that had lost *GAL1* and become 2-DOG^R^. Exposure of cells to each of these genotypic stresses induced significantly elevated frequencies of 2-DOG^R^ colonies (Figure 6D), indicating that DNA damage led to increased chromosome instability in tetraploid cells.

### Evidence that oxidative stress can induce genome instability in tetraploid cells

The experiments described above indicate that endogenous ROS are associated with DNA damage and that this can induce genome instability in tetraploid *C. albicans* cells. We next sought to characterize if exogenous oxidative stress could recapitulate the phenotypes seen on PRE-SPO medium and cause chromosome instability in tetraploid cells. Tetraploid cells were exposed to hydrogen peroxide (H_2_O_2_) as well as the oxidative stress-inducing agents paraquat (PQT) and piperlongumine (PL). PQT generates superoxide anions within the mitochondrial respiratory chain (Cocheme and Murphy, 2008), whereas PL decreases the cellular levels of reduced glutathione thereby promoting ROS accumulation in cells (Raj et al., 2011). When exposed to H_2_O_2_, PQT, or PL for 24 h at 37°C, tetraploid cells stained positively for ROS and Cap1 protein translocated into the nucleus, consistent with cells experiencing increased oxidative stress (Figure 7A,B). A nuclear Gam-GFP signal was also observed in tetraploid cells exposed to each of these agents indicative of DSB formation in the presence of these stressors (Figure 7C,D). Finally, analysis of chromosome stability revealed that H_2_O_2_, PQT, and PL induced elevated levels of *GAL1* marker loss in tetraploid cells (Figure 7E). Together, these results establish that exposure to oxidative stress induces DSBs and chromosome instability in tetraploid cells in a manner that parallels that observed during growth on PRE-SPO medium.

**Figure 7.**
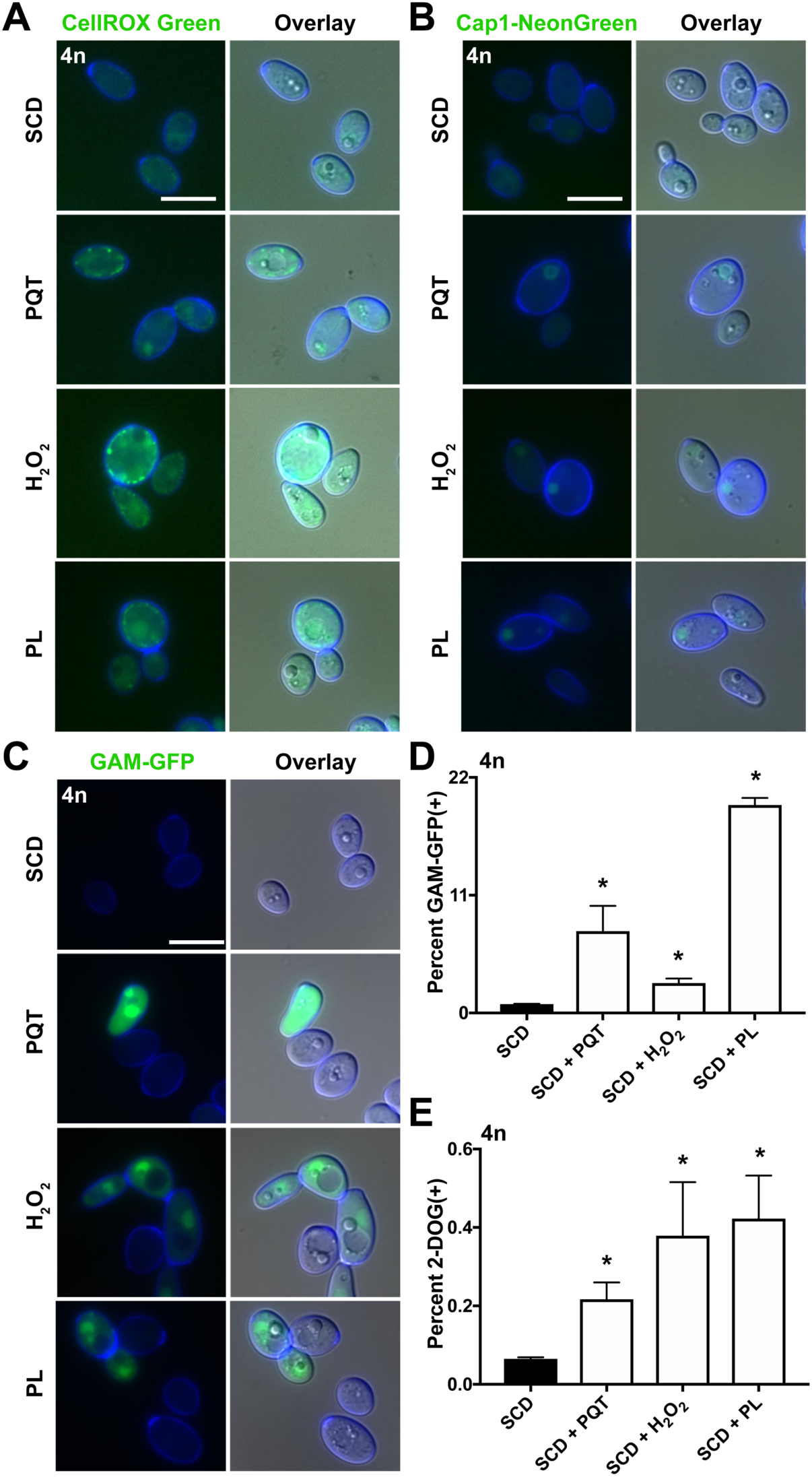
Tetraploid cells exposed to oxidative stress undergo CCL. **(A)** Tetraploid cells were grown on SCD medium supplemented with the oxidative-stress inducing agents paraquat (PQT; 600µg/mL), hydrogen peroxide (H_2_O_2_; 2mM), and piperlongumine (PL; 100µg/mL) at 37°C for 24 h and stained with CellROX Green. Cell images indicate calcofluor white staining (cell wall; blue), GFP, and a merged image of GFP/DAPI/DIC channels. Scale bar = 10 µm. (**B)** Tetraploid cells containing Cap1-mNeonGreen were grown on SCD medium containing PQT (600µg/mL), H_2_O_2_ (2mM), or PL (100µg/mL) at 37°C for 24 h. Cell images as in (A). Scale bar = 10 µm. (**C)** Tetraploid cells expressing a Gam-GFP reporter were cultured on SCD medium supplemented with PQT (600µg/mL), H_2_O_2_ (2mM), or PL (100µg/mL) at 37°C for 24 h and analyzed via fluorescence microscopy. Cell images as in (A). Scale bar = 10 µm. (**D)** Flow cytometric analysis of the tetraploid Gam-GFP reporter strain grown on SCD supplemented with PQT (600µg/mL), H_2_O_2_ (2mM), or PL (100µg/mL) at 37°C for 24 h. (**\***) denotes a significant difference in Gam-GFP values between untreated SCD medium and that supplemented with the indicated agent (p<0.05). (**E)** 2-DOG assay to monitor for loss of *GAL1* function from tetraploid strain RBY18 (*gal1/gal1/GAL1/GAL1*) after culture on SCD medium or SCD medium supplemented with PQT (600µg/mL), H_2_O_2_ (2mM), or PL (100µg/mL) at 37°C for 7 d. (**\***) denotes a significant difference between the percentage of 2-DOG^R^ colonies between cells grown on SCD or SCD supplemented with the indicated agent (p<0.05).

**Figure 8.**
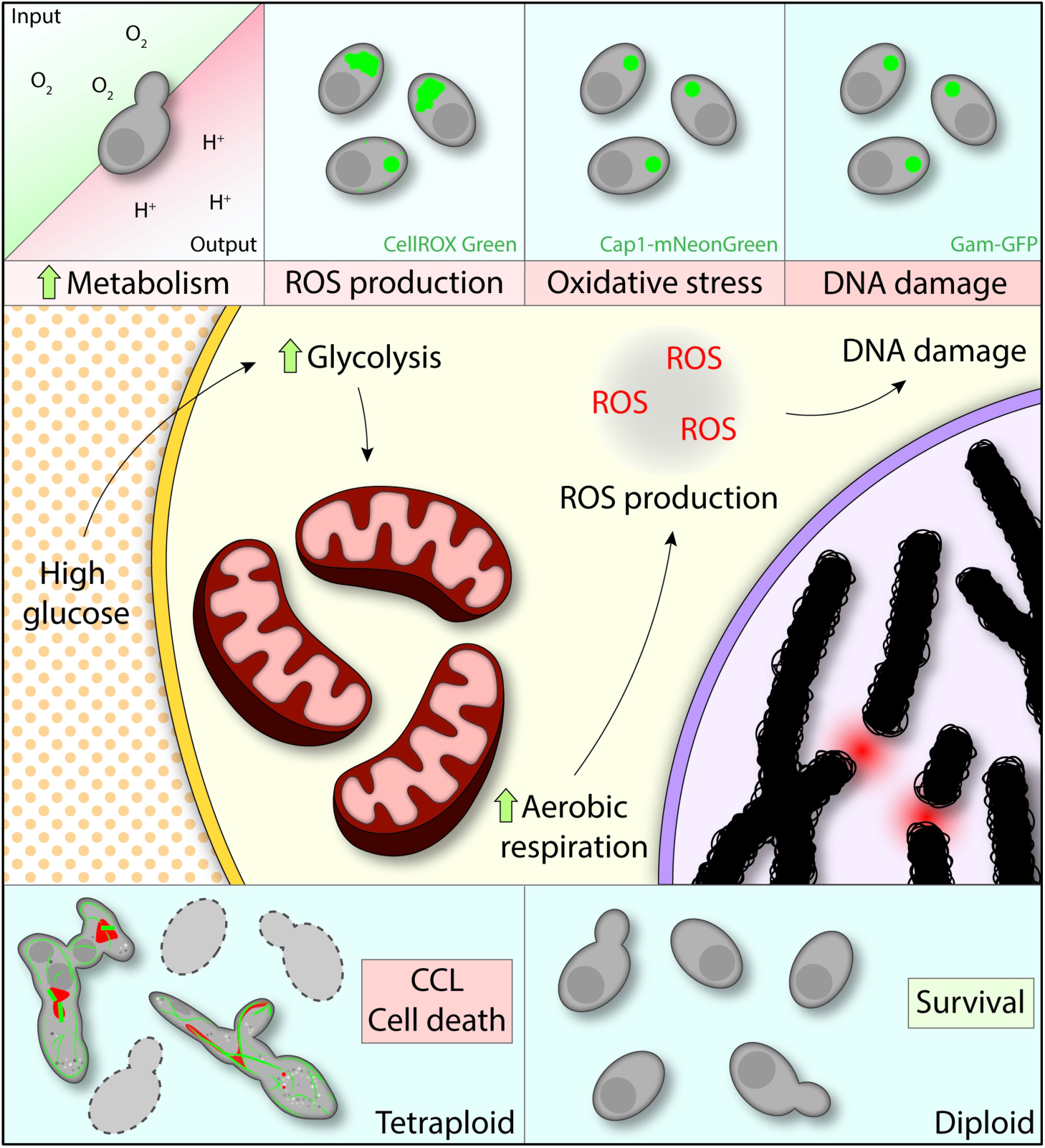
Model of tetraploid cell chromosome instability during culture on PRE-SPO medium. Nutrient concentrations in PRE-SPO medium result in high rates of fermentation and respiration, resulting in ROS production. ROS cause oxidative stress and DNA damage ultimately resulting in cell death and chromosome instability in polyploid cells.

## Discussion

Ploidy change is a natural occurrence in eukaryotic cells being integral to sexual reproduction, the development of certain somatic tissues, and as a driver of tumorigenesis (Bennett et al., 2014; Storchova and Pellman, 2004; Todd et al., 2017). A striking example of ploidy change occurs in *C. albicans* where diploid cells mate to form tetraploid cells that then return to the diploid (or aneuploid) state via aberrant mitotic divisions (Bennett, 2015; Bennett and Johnson, 2003; Forche et al., 2008). Ploidy reduction in tetraploid cells is promoted by growth on glucose-rich PRE-SPO medium whereas diploid cells remain stable under these culture conditions. In this work, we reveal that nutritional cues selectively cause genomic instability and cell death in tetraploid cells and identify causal links to metabolic flux, ROS production, oxidative stress, and DNA double-strand break (DSB) formation. Our results thereby connect metabolic cues to genome instability in polyploid cells and suggest a model by which metabolism regulated ploidy cycling as an evolutionary precursor to a conventional meiosis.

### An elevated metabolism in polyploid cells can enhance ROS levels

*C. albicans* tetraploid cells are stable when cultured on standard yeast media although long-term passaging, as in *S. cerevisiae*, causes cells to converge on the diploid state (Gerstein et al., 2006; Hickman et al., 2015; Selmecki et al., 2015). Growth on PRE-SPO medium accelerates genome instability in a ploidy-specific manner; tetraploid cells show reduced viability and increased chromosomal instability whereas diploid cells do not. Expression profiling revealed only limited differences between diploid and tetraploid *C. albicans* cells, consistent with studies in *S. cerevisiae* where ploidy (and cell size) have a modest, albeit significant, effect on the transcriptome (Galitski et al., 1999; Wu et al., 2010). The more rapid induction of glycolytic and oxidative stress response genes on PRE-SPO than on YPD medium, however, suggested that increased metabolic activity could contribute to the phenotypes observed on PRE-SPO medium.

A direct assessment of *C. albicans* metabolism revealed that PRE-SPO-cultivated cells show increased fermentative and aerobic respiratory rates, indicating that this glucose-rich medium stimulates both metabolic processes. We also found that tetraploid cells exhibit metabolic activities that are 5-6-fold higher than diploid cells, while cell size and mitochondrial content are roughly doubled in tetraploid cells relative to diploid cells. In a recent analysis of mammalian cells mitochondrial content also scaled with cell size, although oxidative phosphorylation was maximal at intermediate cell sizes which could determine the optimal size of mammalian cells (Miettinen and Bjorklund, 2016).

In line with increased metabolic flux and the expression of oxidative stress genes, *C. albicans* cells grown on PRE-SPO (but not YPD) medium showed high levels of ROS production and activation of the ROS-responsive transcription factor Cap1. Supplementation of PRE-SPO medium with antioxidants or overexpression of endogenous antioxidant enzymes suppressed both ROS and the oxidative stress response. The addition of antioxidants may protect against oxidative stress by lowering the overall cellular redox potential as shown in *S. cerevisiae* (Bednarska et al. 2008).

### Endogenous ROS-induced DNA damage and chromosome instability

ROS production is closely linked to DNA damage in multiple cell types (Storr et al., 2013; Tafani et al., 2016) and we observed extensive DSB formation in *C. albicans* cells grown on PRE-SPO medium. Supplementation of this medium with antioxidants mitigated both DSB formation and genetic instability in tetraploid cells. This indicates that ROS generated by a hyperactive metabolism results in DSB formation, which in turn induces genome instability and loss of viability in polyploid cells.

In support of this model, *C. albicans* tetraploid cells were more susceptible to treatment with DNA damaging agents than diploid cells, as evidenced by greater cell death and chromosomal instability when treated with MMS or HU. This result mirrors studies in *S. cerevisiae* where tetraploid cells were found to be more sensitive to DSB-inducing agents than diploid cells and to be more reliant on homologous recombination for DNA repair, particularly at elevated temperatures (Storchova et al., 2006). Cells of higher ploidy are likely to be more susceptible to DNA damage and chromosome mis-segregation due to the altered geometry of the mitotic spindle as cell size increases (Storchova et al., 2006).

We also note that treatment of *C. albicans* tetraploid cells with exogenous oxidative stress-inducing agents recapitulated the phenotypes of tetraploid cells on PRE-SPO medium. Treatment of cells with these agents resulted in elevated levels of ROS, DSB formation, cell death and chromosome instability. This is consistent with studies in *S. cerevisiae* where inactivation of DNA repair genes promoted genome instability due to increased oxidative DNA damage (Degtyareva et al., 2008). The current study also establishes that ROS are particularly consequential in cells of higher ploidy - ROS can induce tetraploid cells to undergo a ploidy reduction to a lower ploidy state.

### Polyploidy and elevated ROS levels in mammalian cells

The results described here in *C. albicans* parallel phenomena observed in mammalian cells during tumorigenesis. Approximately 90% of all solid tumors are aneuploid and a subset of these aneuploid forms arise via transient polyploid intermediates (Coward and Harding, 2014). Polyploid mammalian cells, as in *C. albicans*, exhibit altered metabolism and genome stability relative to their diploid counterparts. For example, polyploid prostate and mammary epithelial cells are marked by increased mitochondrial content (Roh et al., 2012), and this in turn has been implicated in promoting ROS production, aneuploid formation and tumorigenesis (Gibellini et al. 2010). Studies in embryonic fibroblasts and mammary epithelial cells have similarly linked mitochondrial ROS with DNA damage and increased genome instability (Radisky et al., 2005; Samper et al., 2003), and expression of mitochondrial superoxide dismutase can suppress genome instability caused by ROS in thymocytes (van de Wetering et al., 2008). Thus, it is now apparent that metabolic cues are directly implicated in ROS production, DNA damage and chromosomal instability in both fungal and animal cells. This indicates that unicellular eukaryotes can provide mechanistic insights into potentially conserved processes underlying genome integrity and tumorigenesis in polyploid cells.

### A model for metabolism-induced ROS in the evolution of eukaryotic ploidy cycles

A critical unresolved question in biology is how did meiosis and sexual reproduction evolve in eukaryotes? The emergence of sex would have enhanced DNA repair processes and enabled adaptation, yet came with significant costs including the ‘twofold cost’ of sex (Bernstein et al., 2011; de Visser and Elena, 2007; Goddard, 2016; McDonald et al., 2016). Intriguingly, sex may have been particularly advantageous for dealing with oxidative stress, as the endosymbiont that formed the pre-mitochondrion could have generated high levels of endogenous ROS and associated DNA damage (Bernstein et al., 2011; Blackstone, 1995; Horandl and Speijer, 2018). In support of a connection between ROS and sex, oxidative stress promotes sexual reproduction in *Schizosaccharomyces pombe* (Bernstein and Johns, 1989), *Phytophthora cinnamomi* (Reeves and Jackson, 1974), and *Volvox carteri* (Nedelcu et al., 2004), indicating that ROS is a common cue for activation of sexual programs.

The current work suggests how ROS could have facilitated ploidy cycling as a forerunner to meiosis in the early eukaryote. High levels of endogenous ROS (from the pre-mitochondrion) would have generated DNA damage, and this would have been particularly detrimental to genome stability in polyploid cells. Thus, if haploid cells gave rise to diploid or polyploid cells (by cell fusion, endoreduplication, mitotic slippage or a failure in cytokinesis) then the resulting cells would have both generated more ROS and been more susceptible to ROS-induced DNA damage. Higher ploidy cells would therefore have reverted to a more stable, lower ploidy state. In this model, ploidy reduction would have been a natural consequence of the high levels of ROS produced by early eukaryotes (Horandl and Speijer, 2018), and a primitive ploidy cycle would have occurred in the absence of meiosis. Intriguingly, recent studies reveal that metabolic processes can also trigger polyploidization in human cell lines (Tan et al., 2018), indicating that both increases and decreases in ploidy can be facilitated by metabolic activity.

The envisaged model aligns with key aspects of meiosis in extant eukaryotes. DSBs are essential for recombination in meiotic cells with their production tightly regulated by the Spo11 endonuclease in all eukaryotic kingdoms (Lam and Keeney, 2014). Interestingly, however, the role of Spo11 can be bypassed in *C. elegans* by using γ radiation to introduce DSBs (Dernburg et al., 1998), indicating that Spo11 is dispensable if DSBs are generated by an alternative mechanism. It is therefore proposed that ROS-induced DSBs were the causal agents of ploidy reduction in early eukaryotes, and that this step subsequently came under the control of a regulated endonuclease to allow greater precision of potentially lethal DNA breaks.

## Conclusions

In this study, we demonstrate connections between metabolism, endogenous ROS production, DNA damage and genome instability, and establish how polyploid cells can selectively be induced to reduce their ploidy even in the absence of a conventional meiosis. These studies parallel those in cancer cells where polyploidy, ROS and chromosome instability are frequently observed, indicating that similar mechanisms can impact genome stability in diverse eukaryotes. Our results also provide a model for how metabolism-induced processes could have enabled a rudimentary ploidy cycle in eukaryotic evolution. Studies across additional eukaryotic lineages will help to define the relationship between metabolism and genome integrity, and to further infer the steps that led to the complex meiotic program in extant eukaryotes.

## Supporting information

## Acknowledgements

We would like to thank Matthew Anderson and Joshua Wang for assistance with RNA-seq analyses, Marla Tipping and Kate Neville for assistance with Seahorse experiments, Iuliana Ene and Corey Frazer for comments on the manuscript, and Matthew Hurton and Emily Roblee for help with preliminary assays. This work was supported by National Institutes of Health grants AI081704 and AI122011 to RJB and GM110578 and GM 094712 to NA, and by a Burroughs Wellcome Fund PATH award to RJB.

## Materials and Methods

### Media and chemicals

Yeast extract peptone dextrose (YPD) and synthetic complete dextrose (SCD) media were prepared as described (Guthrie and Fink, 1991). *S. cerevisiae* pre-sporulation (“PRE-SPO”) medium contained 0.8% yeast extract, 0.3% peptone, 10% glucose (added prior to autoclaving), and 2% agar. Quercetin hydrate (Acros Organics, #AC174070100), curcumin (Acros Organics, #AC218580100), L-ascorbic acid (Acros Organics, #AC352680050), L-glutathione (Sigma-Aldrich, Cat. No. G4251-10G), dithiothreitol (Fisher Scientific, #BP172-25), paraquat dichloride hydrate (Sigma-Aldrich, #36541-100MG), hydrogen peroxide (Sigma-Aldrich, #H1009-100ML), piperlongumine (Cayman Chemical, #20069-09-4), methyl methanesulfonate (Acros Organics, #AC156890050), and hydroxyurea (MP Biomedicals, #0210202310-10g) were added to the indicated media once the media had cooled to approximately 40°C, at which point the media was poured and plates were used for assays within 24 h.

### Strain construction

A *CAP1*-mNeonGreen cassette was amplified off of pRB895 (*pSFS2A-NeonGreen*) using oligonucleotides 4564 and 4565, which contain homology to the *CAP1* gene and the *mNeonGreen* plasmid. This cassette was transformed into the diploid strain SC5314 and the tetraploid strain RBY18 to create strains CAY9331 (diploid) and CAY9332 (tetraploid).

The plasmid *pRS11* was generating using a *C. albicans* codon-optimized *Gam* gene flanked by two *SalI* restriction sites was synthesized, digested with *SalI*, and cloned into the *pNIM1* vector (Park and Morschhauser, 2005). *pRS11* was linearized with restriction enzymes *SacII* and *ApaI* and transformed into the diploid strain SC5314 and the tetraploid strain RBY18 to generate strains CAY7504 (diploid) and CAY7571 (tetraploid).

The plasmid *pLC605* contains a *Tet-OFF* promoter sequence which can be fused to genes of interest to generate overexpression (O/E) cassettes (Veri et al., 2018). Tet-OFF O/E cassettes were created for the antioxidant enzymes encoded by *SOD2, SOD3, TRX1*, and *GCS1* by PCR amplification off of pLC605 using oligos 4823/4824, 4702/4703, 4813/4814, and 48174818, respectively. These cassettes were transformed into RBY18 to generate strains CAY9516 (*SOD2 O/E*), CAY9388 (*SOD3 O/E*), CAY9501 (*TRX1 O/E*), and CAY9507 (*GCS1 O/E*).

### RNA sequencing

Strains SC5314 and RBY18 were collected from YPD medium, approximately 6 x 10^7^ cells from each strain were top-spread onto YPD and PRE-SPO media and incubated at 30°C and 37°C, respectively. Cells were collected at 3, 6, 12, and 24 h, flash frozen in liquid nitrogen, and stored at −80°C. Each experiment was performed in biological duplicate. RNA was purified using the Ambion RiboPure Yeast RNA Purification Kit (ThermoFisher, #AM1926) and RNA integrity assessed using an Agilent Bioanalyzer 2100. mRNA libraries were prepared using the Illumina TruSeq Stranded mRNA Library Preparation Kit (Illumina, #20020595). Libraries were then pooled and sequenced on an Illumina HiSeq 4000 platform generating 100-bp paired-end reads (at The Ohio State University Comprehensive Cancer Center, Columbus, OH). Reads were aligned to the *C. albicans* SC5314 reference genome (Assembly 21) using the Spliced Transcripts Alignment to a Reference (STAR) aligner (Dobin et al., 2013). Differential expression analysis was conducted using an in-house script that was executed in R. Gene ontology enrichment analysis was conducted using the Gene Ontology Consortium enrichment analysis tool (Mi et al., 2017).

### Viability assays

Strains SC5314 and RBY18 were collected from YPD medium and approximately 6 x 10^7^ cells from each strain were top-spread onto YPD and PRE-SPO media and incubated at 30°C and 37°C, respectively. At 1, 2, 3, 5, and 7 days, cells were collected, stained with 0.5 mg/mL propidium iodide, and imaged using a fluorescence microscope (Zeiss Z1 Axio Observer, excitation filter 555 nm, 250 ms exposures). Images were then viewed with Fiji image processing software and ∼300 cells were scored for viability, with PI-positive cells representing dead yeast cells (Dudgeon et al., 2008). Each experiment was performed in biological quintuplicate.

### 2,3,5-triphenyltetrazolium chloride (TTC) reduction assay

To assess metabolic activity of *C. albicans* cells on YPD and pre-spo media, a TTC overlay technique was used (Ogur et al., 1957). Approximately 2 x 10^7^ cells of SC5314 and RBY18 strains were spotted onto YPD and PRE-SPO media and incubated for 24 h at 30°C and 37°C, respectively. Prior to applying the TTC overlay solution, each plate was imaged for a 0 min reference. A TTC-overlay agar solution (0.067 M potassium phosphate buffer pH 7.0, 1.5% agar, 0.1% TTC) was warmed to 55°C and 10 mL added onto the plates containing the spot-inoculated colonies. The flooded plates were imaged 15 minutes after the addition of the overlay solution.

For image quantitation, collected images were viewed with Fiji image processing software. Spot colonies were selected and mean RGB breakdowns of pixel values for each colony were determined using the color histogram tool. The relative luminance (*Y*) of each colony was calculated by plugging the RGB values into the formula *Y* = 0.2126*R* + 0.7152*G* + 0.0722*B* and the percent of red light contributing to the overall luminance was calculated using the formula (0.2126*R*/*Y*)×100. The percent red light of overall luminance was calculated for both the 0 min and 15 min images, and the 15 min value was normalized to the 0 min value for a final red pixel intensity value.

### Aerobic respiratory and fermentative rate assays

Strains SC5314 and RBY18 were collected from YPD medium and approximately 6 x 10^7^ cells from each strain were top-spread onto YPD and PRE-SPO media and incubated at 37°C for 24 h. An Agilent Seahorse XFe96 sensor cartridge (Agilent Technologies, Cat. No. 102416-100) was hydrated in a utility plate with each well containing 200 µL of sterile water overnight at 37°C. On the following day, the sensor cartridge was moved into a utility plate containing 200 µL of XF Calibrant (Agilent Technologies, Cat. No. 100840-000) in each well and incubated for one hour at 37°C. Cells were harvested from each plate and suspended in sterile water to a concentration of approximately 5.0×10^8^ CFU/mL. 10 µL of each cell suspension was seeded into wells containing 200 µL of pre-warmed Seahorse XF assay medium (supplemented with 10 mM glucose, 10 mM sodium pyruvate) in an Agilent Seahorse XF96 cell culture microplate (Agilent Technologies, #102416-100). The cell plate was placed into the Seahorse XFe96 instrument and the assay was run for seven total cycles, with each cycle consisting of one minute of mixing, zero minutes waiting, and three minutes measuring. Upon completion of assay, the exact number of cells in each cell suspension were assessed via hemocytometer counts and oxygen consumption rate (OCR) and extracellular acidification rate (ECAR) measurements were normalized to both the total number of cells seeded into each well and the viability of cells under each growth condition (calculated in the viability assays detailed above). These normalized measurements were then multiplied by 10^6^ for final OCR measurements in the units of pmol/min/10^6^ CFU and ECAR measurements in the units of mpH/min/10^6^ CFU. Each experiment was performed in biological quintuplicate.

### Antioxidant enzyme overexpression assays

100 µL of 50 µg/mL doxycycline (DOX) was top-spread onto PRE-SPO medium plates (representing a final concentration of ∼2.5 µg/mL doxycycline) and allowed to diffuse into the media. As a control, 100 µL of water was top-spread onto PRE-SPO medium plates. Strains CAY9516 (*SOD2 O/E*), CAY9388 (*SOD3 O/E*), CAY9501 (*TRX1 O/E*), and CAY9507 (*GCS1 O/E*) were collected from YPD medium and approximately 6 x 10^7^ cells from each strain were top-spread onto PRE-SPO medium +DOX and – DOX and incubated for 24 h at 37°C, at which point cells were harvested and their viability was assessed as detailed above.

### Stress assays

SC5314 and RBY18 cells were grown overnight in liquid YPD medium at 30°C. The following day, the cultures were diluted 1:50 and grown to mid-logarithmic phase (approximately 4-5 hours). Cell density was determined by hemocytometer count and cells were diluted to a density of 2×10^8^ CFU/mL. Five successive 1:10 serial dilutions were made in sterile phosphate-buffered saline (PBS) and 5 µL of each cell suspension spotted onto freshly prepared SCD, SCD + 20 mM hydroxyurea, and SCD + 0.01% methyl methanesulfonate media. Plates were incubated for 48 h at 30°C and 37°C at which point plates were imaged.

### CellROX Green staining

SC5314 and RBY18 were streaked into quadrants on YPD and PRE-SPO media and incubated for 24 h at 30°C and 37°C, respectively. Cells were harvested and washed with sterile PBS. PBS-washed cells were then suspended in 5 µM CellROX Green Reagent (ThermoFisher Scientific, #C10444) and incubated for 30 min at 37°C. The staining solution was then removed from cells and cells were washed three times with PBS. CellROX Green-stained cells were then co-stained with calcofluor white at a final concentration of 0.1 mg/mL to visualize the cell wall. For microscopy, cells were imaged using a fluorescence microscope (Zeiss Z1 Axio Observer; CellROX Green: excitation filter 488 nm, 500 ms exposure; Calcofluor white: excitation filter 358 nm, 25 ms exposure) and images viewed and processed with Fiji image processing software. For flow cytometry, a total of 100,000 cells were analyzed on a FACSAria (BD Biosciences) and data was analyzed via FlowJo (TreeStar, Ashland, OR).

### MitoTracker Green staining

SC5314 and RBY18 were streaked into quadrants on YPD and PRE-SPO media and incubated for 24 h at 30°C and 37°C, respectively. Cells were harvested and washed with sterile PBS. PBS-washed cells were then suspended in 1 µM MitoTracker Green Reagent (ThermoFisher Scientific, #M7514) and incubated for 30 min at 37°C. The staining solution was then removed and cells were washed once with PBS. For flow cytometry, a total of 500,000 cells were analyzed on a FACSAria (BD Biosciences) and data was analyzed via FlowJo (TreeStar, Ashaldn, OR).

### Gam-GFP assays

A Gam-GFP reporter strain was used to detect the presence of DSBs in tetraploid *C. albicans* cells. This strain was grown on different media (YPD, PRE-SPO, and PRE-SPO plus antioxidants) in the presence of doxycycline (+DOX; 12.5 µg/ml) to induce the Gam-GFP protein. Plates without doxycycline (-DOX) were included as controls where Gam-GFP was not induced. CAY7504 (diploid) and CAY7571 (tetraploid) cells containing Gam-GFP were inoculated into quadrants on YPD, PRE-SPO, and PRE-SPO plus antioxidants plates and incubated for 24 h at 30°C (YPD) and 37°C (PRE-SPO, PRE-SPO plus antioxidants). Cells were recovered from each plate and suspended in 1 mL of sterile PBS. A total of 500,000 cells from each sample were run on a BD FACSAria (Becton Dickinson Biosciences) and data analyzed using FlowJo software (Tree Star, Ashland, OR). For microscopy, cells were co-stained with calcofluor white at a final concentration of 0.1 mg/mL. Cells were imaged using a fluorescence microscope (Zeiss Z1 Axio Observer; Gam-GFP: excitation filter 488 nm, 250 ms exposure; Calcofluor white: excitation filter 358 nm, 25 ms exposures) and images were viewed and processed with Fiji image processing software.

### Cap1-mNeonGreen assays

Strains CAY9331 (diploid) and CAY9332 (tetraploid) expressing Cap1-mNeonGreen were inoculated into quadrants on YPD, PRE-SPO, and PRE-SPO plus antioxidants media and incubated for 24 h at 30°C (YPD) and 37°C (pre-spo, pre-spo plus antioxidants). Cells were recovered from each plate and suspended in sterile PBS. Cells were co-stained with calcofluor white at a final concentration of 0.1 mg/mL. Cells were imaged using a fluorescence microscope (Zeiss Z1 Axio Observer; CAP1-NeonGreen: excitation filter 488 nm, 250 ms exposures; Calcofluor white: excitation filter 358 nm, 25 ms exposure) and images were viewed and processed with Fiji image processing software.

### Chromosome loss quantification using 2-DOG

RBY18 was streaked out into quadrants on media of interest and incubated for 7 days at 37°C. Cells were then scraped off each plate and suspended in sterile PBS. Two successive 1:100 dilutions were made in sterile PBS. 100 µL of the undiluted and 1:100 dilution cell suspensions were plated onto SC 2-DOG medium and 100 µL of the 1:100 dilution and 1:10000 dilution cell suspensions were plated onto YPD medium. These plates were incubated at 30°C for 48 h, at which point the plates were collected and the total number of colonies were counted on each plate.

### DNA staining for ploidy analysis

SC5314, RBY18, and 2-DOG^R^ colonies were grown overnight at 30°C in YPD medium, at which point cells were collected, fixed with 70% ethanol, and incubated for 1 h at 4°C. Cells were then washed twice with 50 mM TrisCl pH 8.0, 5 mM EDTA, suspended in 2 mg/mL RNase A (Sigma-Aldrich, #R5503-500MG) in 50 mM TrisCl pH 8.0, 5 mM EDTA and incubated for 2 h at 37°C. RNase-treated cells were pelleted and resuspended in 5 mg/mL pepsin (Sigma-Aldrich, #P7000-500G) in 55 mM HCl and incubated for 1 h at 37°C. Cells were then washed twice with 50 mM TrisCl pH 7.5, 5 mM EDTA and suspended in 1 µM SYTOX Green Nucleic Acid Stain (Thermo Fisher Scientific, #S7020) in 50 mM TrisCL pH 7.5, 5 mM EDTA and incubated overnight at 4°C. For ploidy analysis, 100,000 cells were run on an Attune NxT Flow Cytometer (Thermo Fisher Scientific) and data was analyzed via FlowJo (TreeStar, Ashaldn, OR).

## Supplementary Figures

**Supplemental Figure S1.**
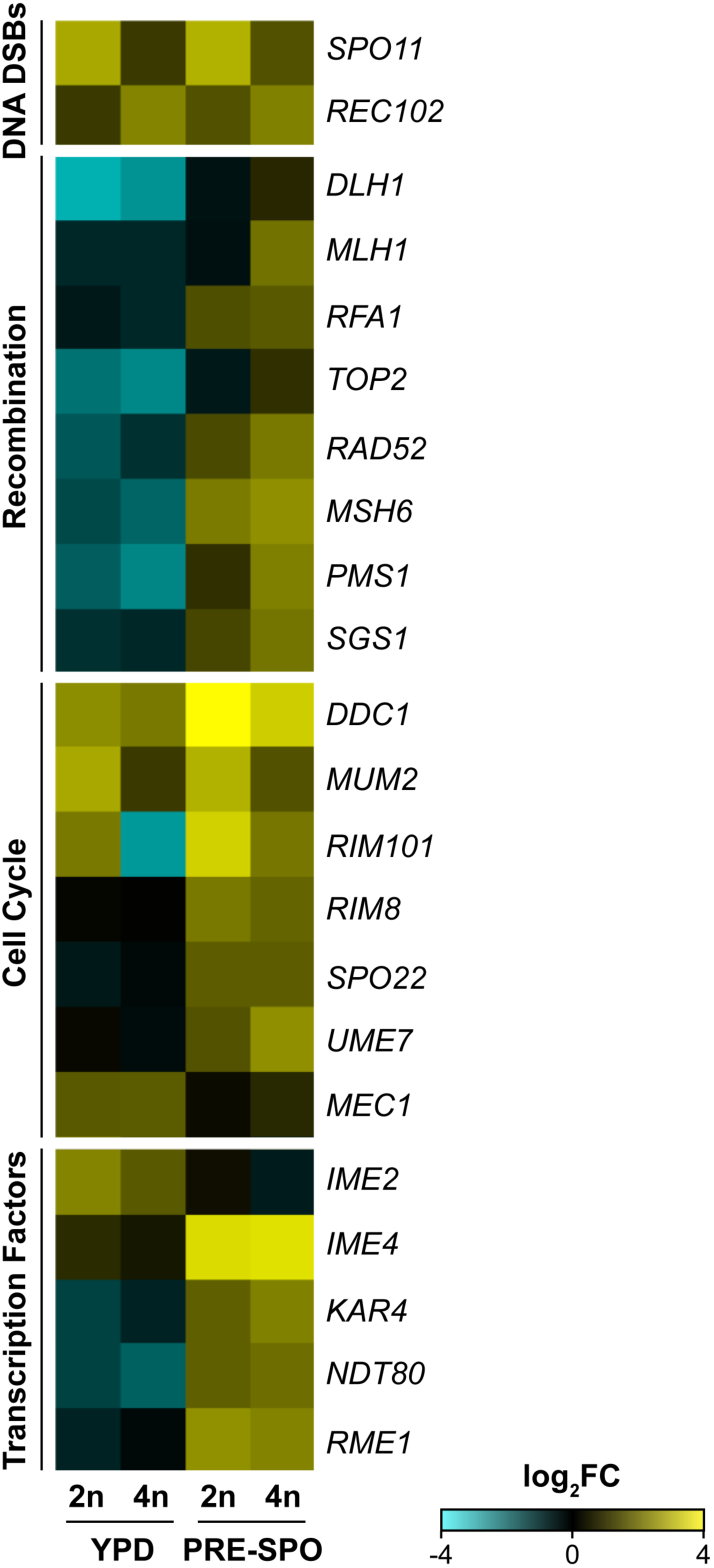
Transcriptional profiling of meiosis genes on YPD and PRE-SPO media. Heatmap of RNA-Seq data for *C. albicans* homologs to “meiotic” genes in diploid and tetraploid cells on YPD and PRE-SPO media at 24 h, including programmed formation of DNA double-strand breaks (DSBs), recombination, coordination of the meiotic cell cycle, and transcription factors.

**Supplemental Figure S2.**
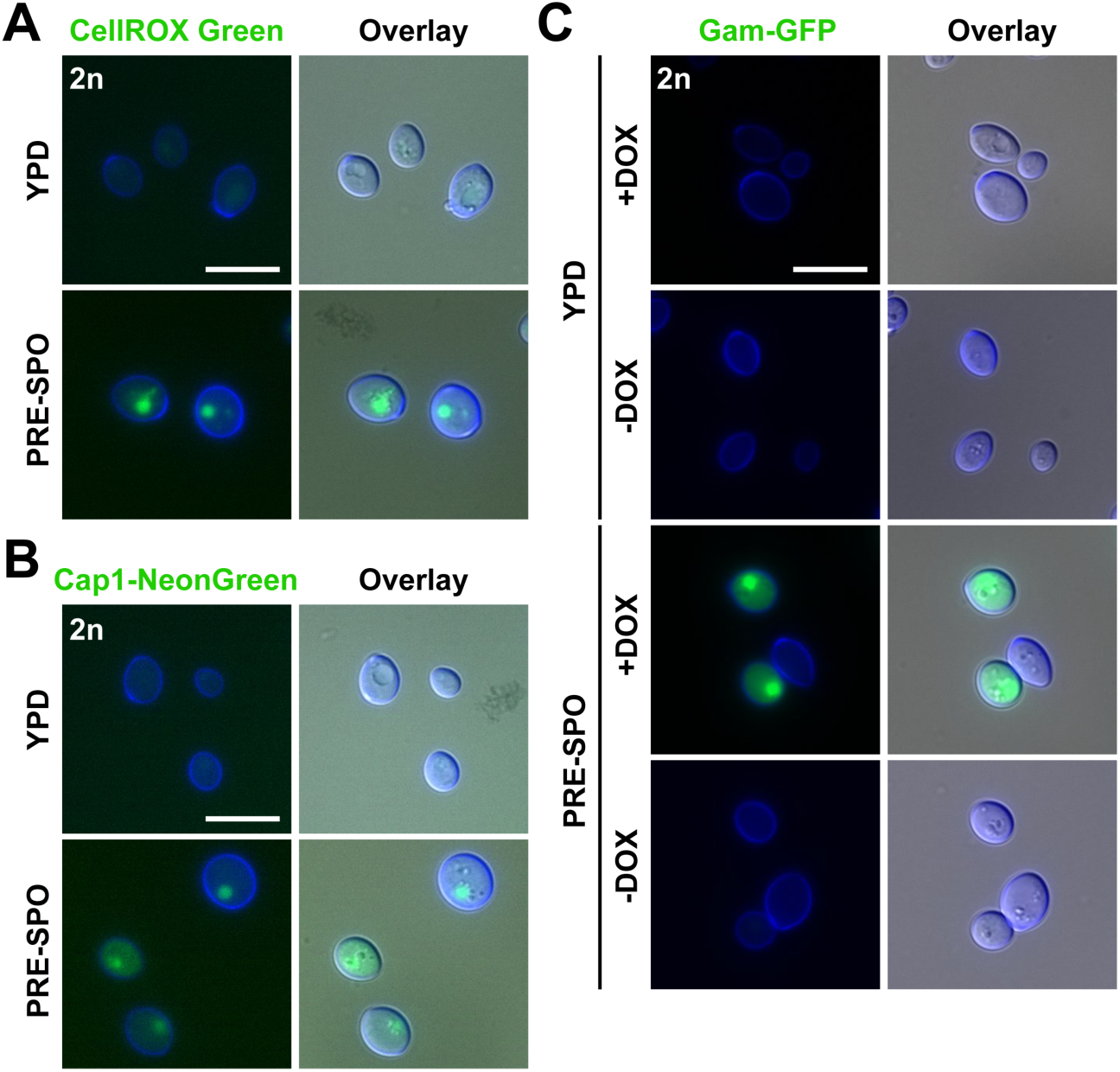
Diploid response to growth on PRE-SPO medium. **(A)** Diploid cells were grown on YPD or PRE-SPO medium and stained with CellROX Green. Cell images indicate calcofluor white staining (cell wall; blue), GFP, and a merged image of GFP/DAPI/DIC channels. Scale bar = 10 µm. (**B)** Diploid cells containing a fluorescently labeled version of the transcription factor Cap1 (Cap1-mNeonGreen) were grown on YPD or PRE-SPO medium at 30°C or 37°C, respectively, for 24 h. Cell images indicate calcofluor white staining (cell wall; blue), GFP, and a merged image of GFP/DAPI/DIC channels. Scale bar = 10 µm. (**C)** A diploid strain expressing a Tet-ON Gam-GFP reporter was cultured on YPD or PRE-SPO medium at 30°C or 37°C, respectively, in the presence (Gam-GFP ON) or absence (Gam-GFP OFF) of doxycycline for 24 h. Cell images indicate calcofluor white staining (cell wall; blue), GFP, and a merged image of GFP/DAPI/DIC channels. Scale bar = 10 µm.

**Supplemental Figure S3.**
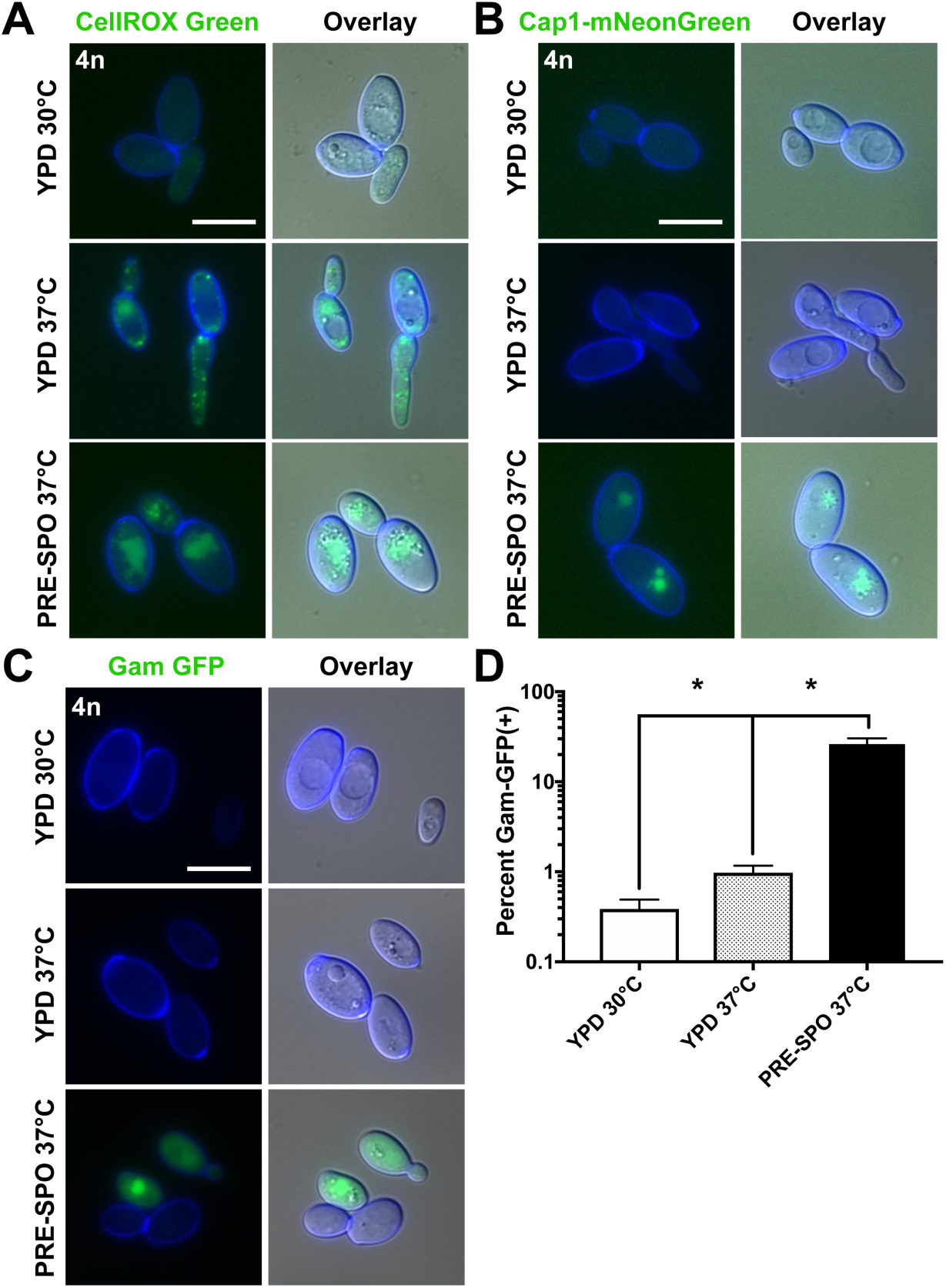
Tetraploid response to growth on YPD medium at 37°C. **(A)** Tetraploid cells were grown on YPD medium at 30°C or 37°C, or PRE-SPO medium at 37°C, for 24 h and stained with CellROX Green. Cell images indicate calcofluor white staining (cell wall; blue), GFP, and a merged image of GFP/DAPI/DIC channels. Scale bar = 10 µm. (**B)** Tetraploid cells containing a fluorescently labeled version of the transcription factor Cap1 (Cap1-mNeonGreen) were grown on YPD medium at 30°C or 37°C, or PRE-SPO medium at 37°C, for 24 h. Cell images indicate calcofluor white staining (cell wall; blue), GFP, and a merged image of GFP/DAPI/DIC channels. Scale bar = 10 µm. (**C)** A diploid strain expressing a Tet-ON Gam-GFP reporter was cultured on YPD medium at 30°C or 37°C, or PRE-SPO medium at 37°C, in the presence (Gam-GFP ON) of doxycycline for 24 h. Cell images indicate calcofluor white staining (cell wall; blue), GFP, and a merged image of GFP/DAPI/DIC channels. Scale bar = 10 µm. (**D)** Flow cytometric analysis of the tetraploid Gam-GFP reporter strain grown on YPD medium at 30°C or 37°C, or PRE-SPO medium at 37°C in the presence of doxycycline for 24 h. (**\***) indicates a significant difference (p<0.05).

**Supplemental Figure S4.**
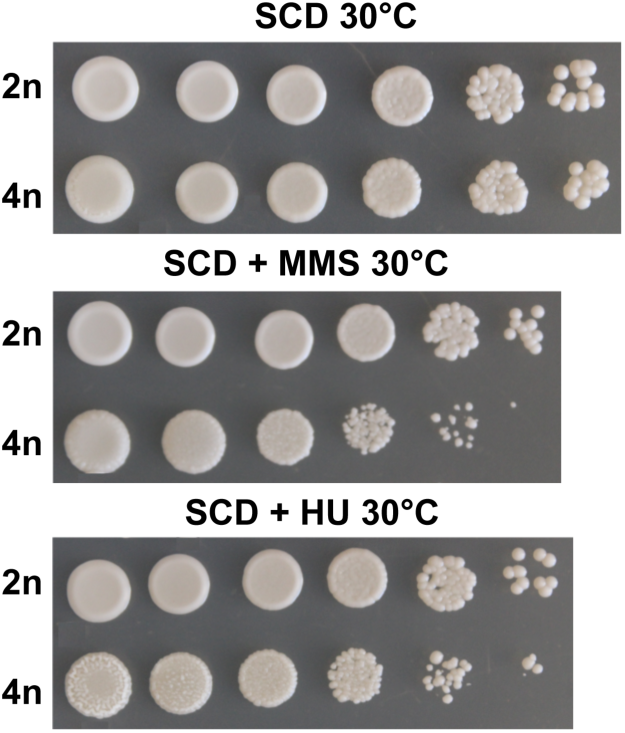
MMS and HU stress assay at 30°C. 10-fold dilutions of diploid and tetraploid cells were spot inoculated onto SCD medium and SCD medium containing the DNA-damaging agents HU and MMS. Cells were allowed to grow for 48 h at 30°C and plates imaged.

**Supplemental Table S1.**
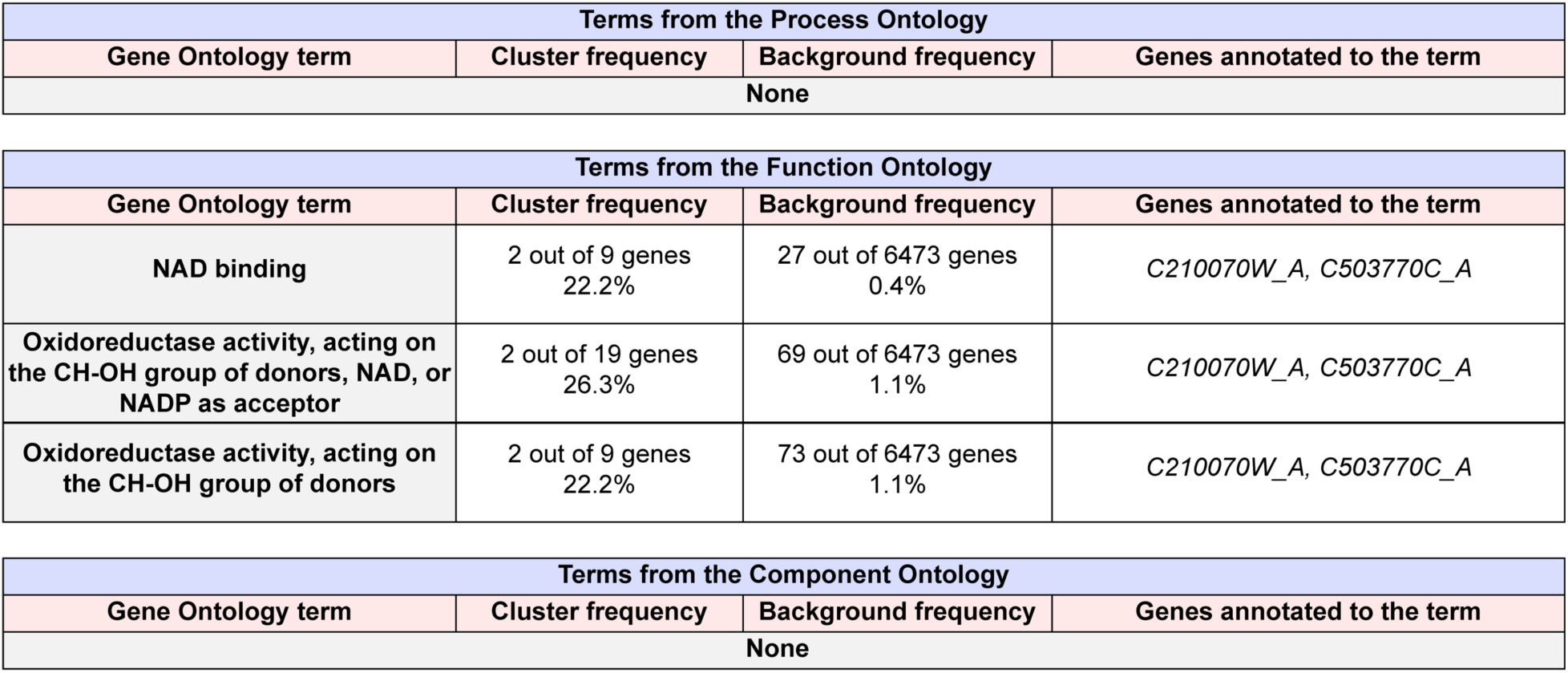
Results of gene ontology (GO) term enrichment analysis of differentially expressed genes between diploid and tetraploid cells on PRE-SPO medium at 3 h. Background frequency represents the total number of genes annotates to each GO term in the *C. albicans* genome divided by the total number of genes in the genome. All included terms are significantly enriched (p<0.05).

**Supplemental Table S2.** Results of GO term enrichment analysis of differentially expressed genes between diploid and tetraploid cells on PRE-SPO medium at 6 h. Background frequency represents the total number of genes annotates to each GO term in the *C. albicans* genome divided by the total number of genes in the genome. All included terms are significantly enriched (p<0.05).

| Terms from the Process Ontology |  |  |  |
| --- | --- | --- | --- |
| Gene Ontology term | Cluster frequency | Background frequency | Genes annotated to the term |
| Acetate catabolic process | 2 out of 19 genes<br>10.5% | 8 out of 6473 genes<br>0.1% | *CTN1, ICL1* |
| Regulation of flocculation | 2 out of 19 genes<br>10.5% | 9 out of 6473 genes<br>0.1% | *ALS1, PGA13* |
| Acetate metabolic process | 2 out of 19 genes<br>10.5% | 13 out of 6473 genes<br>0.1% | *CTN1, ICL1* |

| Terms from the Function Ontology |  |  |  |
| --- | --- | --- | --- |
| Gene Ontology term | Cluster frequency | Background frequency | Genes annotated to the term |
| Structural constituent of ribosome | 5 out of 19 genes<br>26.3% | 162 out of 6473 genes<br>2.5% | *RDN18, RDN25, RDN5, RDN58, RRNS* |
| Structural molecule activity | 5 out of 19 genes<br>26.3% | 261 out of 6473 genes<br>4.0% | *RDN18, RDN25, RDN5, RDN58, RRNS* |

| Terms from the Component Ontology |  |  |  |
| --- | --- | --- | --- |
| Gene Ontology term | Cluster frequency | Background frequency | Genes annotated to the term |
| Ribosomal subunit | 5 out of 19 genes<br>26.3% | 171 out of 6473 genes<br>2.6% | *RDN18, RDN25, RDN5, RDN58, RRNS* |
| Cytosolic ribosome | 4 out of 19 genes<br>21.1% | 101 out of 6473 genes<br>1.6% | *RDN18, RDN25, RDN5, RDN58* |
| Ribosome | 5 out of 19 genes<br>26.3% | 225 out of 6473 genes<br>3.5% | *RDN18, RDN25, RDN5, RDN58, RRNS* |
| Cytosolic large ribosomal subunit | 3 out of 19 genes<br>15.8% | 58 out of 6473 genes<br>0.9% | *RDN25, RDN5, RDN58* |
| Cytosolic part | 4 out of 19 genes<br>21.1% | 154 out of 6473 genes<br>2.4% | *RDN18, RDN25, RDN5, RDN58* |
| Cell wall | 4 out of 19 genes<br>21.1% | 163 out of 6473 genes<br>2.5% | *ALS1, HWP1, PGA13, PGA31* |
| Fungal-type cell wall | 4 out of 19 genes<br>21.1% | 163 out of 6473 genes<br>2.5% | *ALS1, HWP1, PGA13, PGA31* |
| External encapsulating structure | 4 out of 19 genes<br>21.1% | 164 out of 6473 genes<br>2.5% | *ALS1, HWP1, PGA13, PGA31* |
| Cell surface | 4 out of 19 genes<br>21.1% | 203 out of 6473 genes<br>3.1% | *ALS1, HWP1, PGA13, PGA31* |

**Supplemental Table S3.** Results of gene ontology (GO) term enrichment analysis of differentially expressed genes between diploid and tetraploid cells on PRE-SPO medium at 3 h. Background frequency represents the total number of genes annotates to each GO term in the *C. albicans* genome divided by the total number of genes in the genome. All included terms are significantly enriched (p<0.05).

| Terms from the Process Ontology |  |  |  |
| --- | --- | --- | --- |
| Gene Ontology term | Cluster frequency | Background frequency | Genes annotated to the term |
| None |  |  |  |

| Terms from the Function Ontology |  |  |  |
| --- | --- | --- | --- |
| Gene Ontology term | Cluster frequency | Background frequency | Genes annotated to the term |
| None |  |  |  |

| Terms from the Component Ontology |  |  |  |
| --- | --- | --- | --- |
| Gene Ontology term | Cluster frequency | Background frequency | Genes annotated to the term |
| Chromosomal part | 7 out of 27 genes<br>25.9% | 384 out of 6473 genes<br>5.9% | *CDC6, HCM1, HHT2, MCD1, OGG1, PRI2, TOS4* |
| Chromosome | 7 out of 27 genes<br>25.9% | 404 out of 6473 genes<br>6.2% | *CDC6, HCM1, HHT2, MCD1, OGG1, PRI2, TOS4* |
| Nuclear chromosome part | 6 out of 27 genes<br>22.2% | 311 out of 6473 genes<br>4.8% | *CDC6, HCM1, HHT2, MCD1, PRI2, TOS4* |
| Protein-DNA complex | 3 out of 27 genes<br>11.1% | 56 out of 6473 genes<br>0.9% | *CDC6, HHT2, PRI2* |

**Supplemental Table S4.**
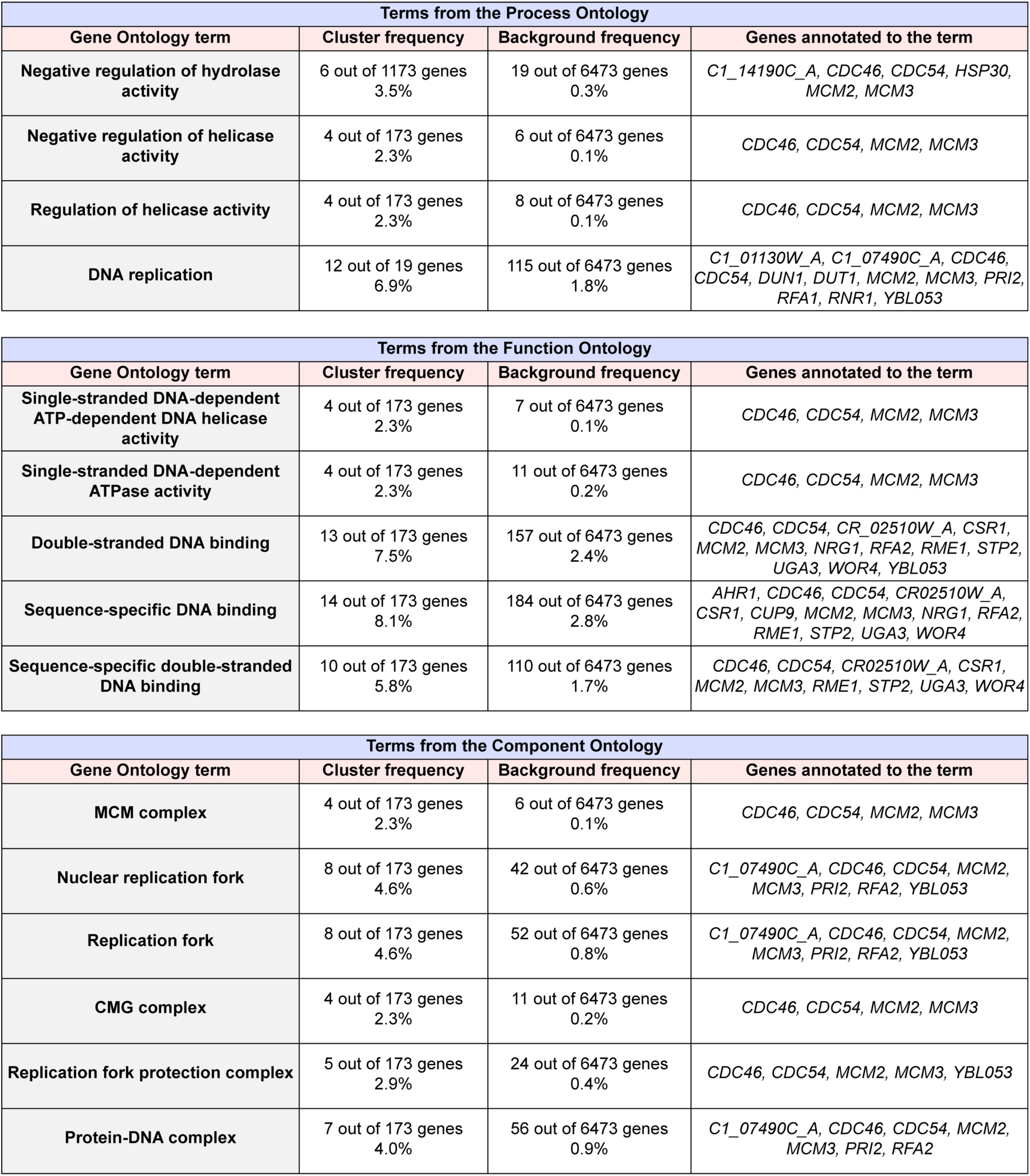
Results of gene ontology (GO) term enrichment analysis of differentially expressed genes between diploid and tetraploid cells on PRE-SPO medium at 24 h. Background frequency represents the total number of genes annotates to each GO term in the *C. albicans* genome divided by the total number of genes in the genome. All included terms are significantly enriched (p<0.05).

| Terms from the Process Ontology |  |  |  |
| --- | --- | --- | --- |
| Gene Ontology term | Cluster frequency | Background frequency | Genes annotated to the term |
| Negative regulation of hydrolase activity | 6 out of 1173 genes<br>3.5% | 19 out of 6473 genes<br>0.3% | *C1_14190C_A, CDC46, CDC54, HSP30, MCM2, MCM3* |
| Negative regulation of helicase activity | 4 out of 173 genes<br>2.3% | 6 out of 6473 genes<br>0.1% | *CDC46, CDC54, MCM2, MCM3* |
| Regulation of helicase activity | 4 out of 173 genes<br>2.3% | 8 out of 6473 genes<br>0.1% | *CDC46, CDC54, MCM2, MCM3* |
| DNA replication | 12 out of 19 genes<br>6.9% | 115 out of 6473 genes<br>1.8% | *C1_01130W_A, C1_07490C_A, CDC46, CDC54, DUN1, DUT1, MCM2, MCM3, PRI2, RFA1, RNR1, YBL053* |

| Terms from the Function Ontology |  |  |  |
| --- | --- | --- | --- |
| Gene Ontology term | Cluster frequency | Background frequency | Genes annotated to the term |
| Single-stranded DNA-dependent ATP-dependent DNA helicase activity | 4 out of 173 genes<br>2.3% | 7 out of 6473 genes<br>0.1% | *CDC46, CDC54, MCM2, MCM3* |
| Single-stranded DNA-dependent ATPase activity | 4 out of 173 genes<br>2.3% | 11 out of 6473 genes<br>0.2% | *CDC46, CDC54, MCM2, MCM3* |
| Double-stranded DNA binding | 13 out of 173 genes<br>7.5% | 157 out of 6473 genes<br>2.4% | *CDC46, CDC54, CR_02510W_A, CSR1, MCM2, MCM3, NRG1, RFA2, RME1, STP2, UGA3, WOR4, YBL053* |
| Sequence-specific DNA binding | 14 out of 173 genes<br>8.1% | 184 out of 6473 genes<br>2.8% | *AHR1, CDC46, CDC54, CR02510W_A, CSR1, CUP9, MCM2, MCM3, NRG1, RFA2, RME1, STP2, UGA3, WOR4* |
| Sequence-specific double-stranded DNA binding | 10 out of 173 genes<br>5.8% | 110 out of 6473 genes<br>1.7% | *CDC46, CDC54, CR02510W_A, CSR1, MCM2, MCM3, RME1, STP2, UGA3, WOR4* |

| Terms from the Component Ontology |  |  |  |
| --- | --- | --- | --- |
| Gene Ontology term | Cluster frequency | Background frequency | Genes annotated to the term |
| MCM complex | 4 out of 173 genes<br>2.3% | 6 out of 6473 genes<br>0.1% | *CDC46, CDC54, MCM2, MCM3* |
| Nuclear replication fork | 8 out of 173 genes<br>4.6% | 42 out of 6473 genes<br>0.6% | *C1_07490C_A, CDC46, CDC54, MCM2, MCM3, PRI2, RFA2, YBL053* |
| Replication fork | 8 out of 173 genes<br>4.6% | 52 out of 6473 genes<br>0.8% | *C1_07490C_A, CDC46, CDC54, MCM2, MCM3, PRI2, RFA2, YBL053* |
| CMG complex | 4 out of 173 genes<br>2.3% | 11 out of 6473 genes<br>0.2% | *CDC46, CDC54, MCM2, MCM3* |
| Replication fork protection complex | 5 out of 173 genes<br>2.9% | 24 out of 6473 genes<br>0.4% | *CDC46, CDC54, MCM2, MCM3, YBL053* |
| Protein-DNA complex | 7 out of 173 genes<br>4.0% | 56 out of 6473 genes<br>0.9% | *C1_07490C_A, CDC46, CDC54, MCM2, MCM3, PRI2, RFA2* |

